# The structure of the F_420_-dependent sulfite-detoxifying enzyme from *Methanococcales* reveals a prototypical sulfite-reductase with assimilatory traits

**DOI:** 10.1101/2022.04.07.487323

**Authors:** Marion Jespersen, Antonio J. Pierik, Tristan Wagner

## Abstract

The coenzyme F_420_-dependent sulfite reductase (Fsr group I) protects hydrogenotrophic methanogens, one of the main contributors in worldwide methane emission, from toxic sulfite. Fsr is a single peptide composed of a F_420_H_2_-oxidase and a novel class of sulfite reductase. Both catalytic domains have been proposed to be the ancestors of modern F_420_-oxido/reductases and dissimilatory/assimilatory sulfite reductases. Here, we describe the X-ray crystal structures of Fsr natively isolated from *Methanocaldococcus jannaschii* (*Mj*Fsr) and *Methanothermococcus thermolithotrophicus* (*Mt*Fsr), respectively refined to 2.30 Å and 1.55 Å resolution. In both organisms, Fsr oligomerizes as a 280-kDa homotetramer, where each siroheme–[4Fe–4S] is catalytically active, in contrast to dissimilatory homologues. The siroheme–[4Fe–4S], embedded in the sulfite reductase domain, is electronically connected to the flavin in the F_420_H_2_-oxidase domain by five [4Fe–4S]-clusters. EPR spectroscopy determined the redox potentials of these [4Fe–4S]^2+/1+^ clusters (−435 to -275 mV), through which electrons flow from FAD to the siroheme–[4Fe–4S]^2+/1+^ (siroheme, -114 mV; [4Fe–4S] -445 mV). The electron relay is mainly organized by two inserted ferredoxin modules, which stabilize the higher degree of oligomerization. While the F_420_H_2_-oxidase part is similar to the β-subunit of F_420_-reducing hydrogenases, the sulfite reductase domain is structurally analogous to dissimilatory sulfite reductases, whereas its siroheme–[4Fe–4S] cofactor is bound in the same way as in assimilatory ones. Accordingly, the reaction of *Mt*Fsr is unidirectional, reducing sulfite or nitrite with F_420_H_2_. Our results provide the first structural insights into this unique fusion, a snapshot of a primitive sulfite reductase that turns a poison into an elementary block of Life.

## Introduction

When cold seawater permeates through sediments or enters hydrothermal vent walls, a partial oxidation of sulfide (HS^-^, S^2-^) results in the formation of (bi)sulfite (HSO_3_^-^), SO_3_^2-^, a highly reactive intermediate of the sulfur cycle (1). Methanogenic archaea are extremely sensitive to this strong nucleophile. It is known to inactivate methanogenesis, which results in the collapse of their central energy metabolism (2, 3). Despite its toxic effect, many hydrogenotrophic methanogens thrive in environments where they are exposed to fluctuating SO_3_^2-^ concentrations, especially methanogens living close to hydrothermal vents or geothermally heated sea sediments (4, 5).

The hyperthermophile *Methanocaldococcus jannaschii*, isolated from a hydrothermal vent (6), expresses the coenzyme F_420_-dependent sulfite reductase (Fsr) in the presence of SO_3_^2-^ (4). This enzyme confers a resistance of up to 40 mM SO_3_^2-^ and allows the methanogen to grow by assimilating SO_3_^2-^ as a sole sulfur source (e.g., absence of S^2-^, (4, 7)). Due to this trait, the *fsr* gene has been used as a genetic marker (7, 8).

Two groups of Fsr were distinguished by phylogenetic analyses (5, 9). While Group II Fsr was found in the genomes of anaerobic methane oxidizers (ANME) and *Methanosarcinales*, Group I Fsr is mostly present in methanogens that lack cytochromes, such as *M. jannaschii*, which belongs to the order of *Methanococcales*. In contrast to the functionally characterized Group I, the physiological role of Fsr Group II is still unknown. Fsr Group II is claimed to be unrelated to SO_3_^2-^ detoxification since 1 mM SO_3_^2-^ inhibits growth of ANME and *Methanococcoides burtonii* (5).

Fsr is organized into two main modules. Its N-terminal half belongs to the F_420_-reducing hydrogenase β-subunit (FrhB) family, together with the F_420_H_2_:quinone oxidoreductase (FqoF), F_420_H_2_:phenazine oxidoreductase (FpoF), putative F_420_-dependent glutamate synthase and formate dehydrogenase (GOGAT and FdhB). Its C-terminal half is composed of a single sulfite/nitrite reductase repeat ((10), S/NiRR, from here on referred to as sulfite reductase domain).

All known sulfite reductases reduce SO_3_^2-^ by using a magnetically coupled siroheme–cysteine–[4Fe–4S] center (11). This specialized cofactor is also used by nitrite reductases to reduce nitrite, a side reaction observed in many sulfite reductases (12). Until now, several groups of sulfite reductases have been identified which are generally classified into assimilatory (aSir, Group I Fsr and presumably alSir/Group I Dsr-like proteins (Dsr-LP)) or dissimilatory (DsrA, DsrB and AsrC) ones depending on their biological function, spectroscopic properties, and molecular composition (12–14). Additionally, there are two biochemically uncharacterized predicted sulfite reductases, Group III Dsr-LP and Group II Fsr (5, 9). The C-terminal half of Fsr is phylogenetically distant from aSirs (here, aSirs refer to siroheme-containing protein subunit) and dSirs (here, dSirs refer to DsrAB) but closely related to Group III Dsr-LP, which are abundant in methanogens (4, 9). However, so far, the only structural data was obtained from aSirs and dSirs, therefore this study will mainly focus on the comparison with those. While aSirs are monomeric enzymes that reduce SO_3_^2-^ directly to S^2-^, to incorporate the latter into biomass, dissimilatory enzymes are organized by the heterodimers DsrA/DsrB, in which DsrA harbors an inactive catalytic site (referred as structural, Fig. S1 (12, 13, 15, 16)). Under physiological conditions, dSirs catalyze the first two-electron reduction step. For the complete reduction of SO_3_^2-^ to S^2-^, dSirs additionally require the sulfur-carrier protein DsrC as well as a membrane complex, which catalyzes the final four-electron reduction for energy conservation (DsrMKJOP complex in *Desulfovibrio vulgaris*, Fig. S1, (17)). DsrAB by itself releases some S^2-^ as well as the reaction intermediates trithionate and thiosulfate (17, 18).

Structural and evolutionary studies suggest that aSirs and dSirs originated from a common progenitor (13, 16), a primitive Sir which harbored a catalytic siroheme–[4Fe–4S] and was operating by itself. The gene coding for the ancestral enzyme was duplicated, and in the dSir case, the duplicated version evolved into DsrB, while DsrA was retained for a structural function. In the case of aSir, the original and duplicated genes fused and only one active siroheme–[4Fe–4S] was retained. Based on sequence and phylogenetic analyses, it has been suggested that *fsr* evolved prior to the duplication event and therefore represents a primordial sulfite reductase in comparison to aSirs and dSirs (4, 9, 19). Alternatively, *fsr* could have arisen through lateral gene transfer followed by gene fusion events.

Besides the evolutionary importance, Fsr is the only known sulfite reductase directly fused to its electron donor module, allowing it to perform the whole SO_3_^2-^ reduction reaction without an additional partner. Fsr reacts with reduced F_420_ (F_420_H_2_), a deazaflavin derivative highlighted as a common electron carrier across many species and is present at high cytoplasmic concentrations in methanogens (4, 20–22). Hydrogenotrophic methanogens mainly reduce F_420_ by the F_420_-reducing hydrogenase (Frh) that performs the two-electron (hydride) transfer via a flavin harbored in its β-subunit (FrhB (23–25)). In Fsr, the oxidation of reduced F_420_ should operate in its N-terminal FrhB-like domain. The electrons are transmitted to the siroheme–[4Fe–4S] by an unknown path that requires a [Fe–S]-cluster relay. These clusters are contained in two ferredoxin domains, the first one located at the N-terminal half, and the second one is inserted in the sulfite reductase domain (Fig. 1). Because of the difference in the redox-potential of the F_420_/F_420_H_2_ (Δ*E*^0^′ = −350 mV) and SO_3_^2-^/S^2-^ (Δ*E*^0′^ = −116 mV) couples, the overall reaction is extremely exergonic (Δ*G*^0′^ = −135 kJ.mol^-1^ per converted HSO_3_^-^) promoting SO_3_^2-^ detoxification at very high rates (4). Its hyperthermophilicity and natural efficiency highlight Fsr as a robust and interesting catalyst for chemists.

**Figure 1.**
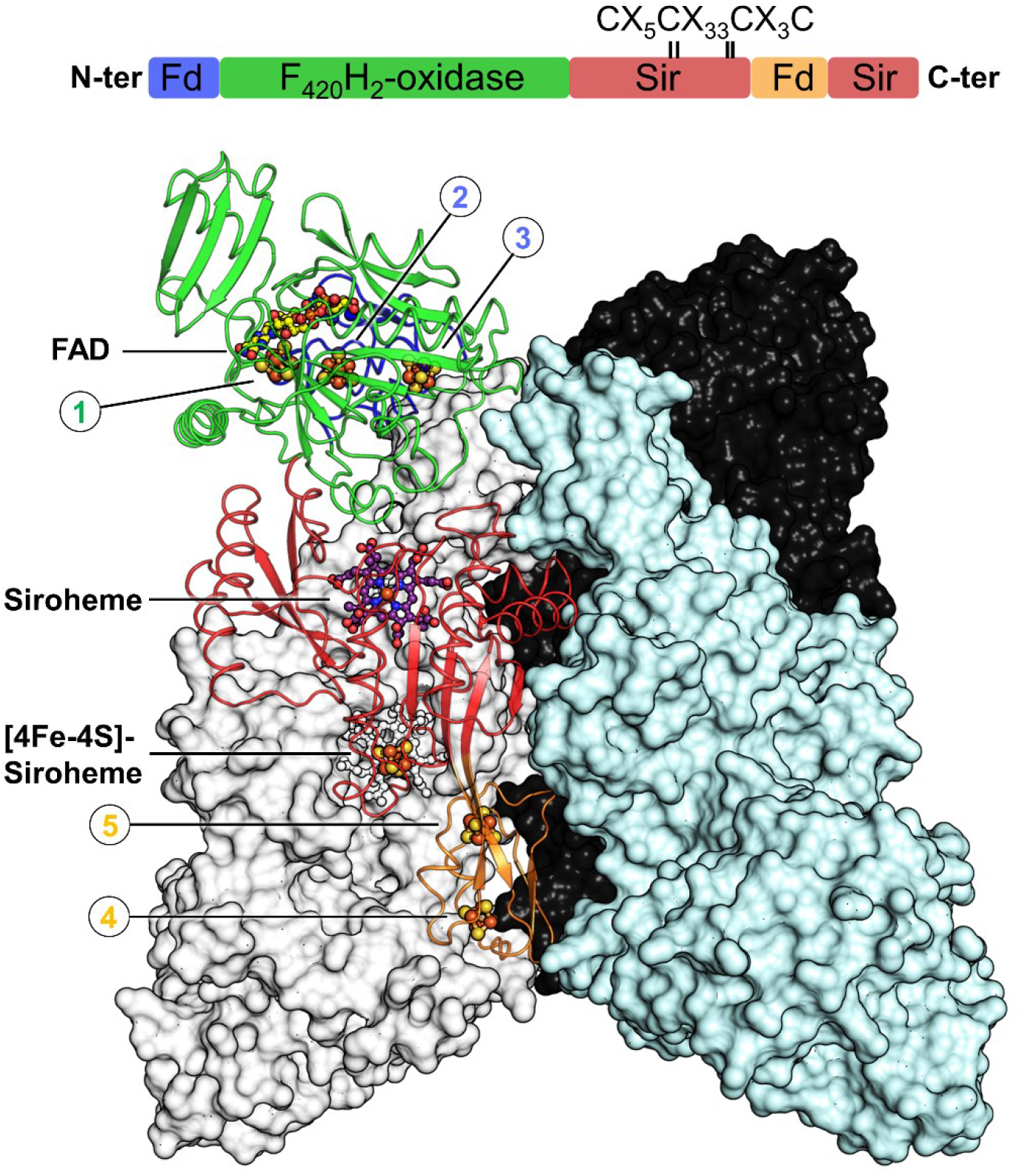
Domain and structural organization of *Mt*Fsr. Visualization of *Mt*Fsr domains (top panel). The [4Fe–4S]-cluster binding motif in the proximity of the siroheme is highlighted. The main panel shows the tetrameric arrangement of *Mt*Fsr. Three chains are represented in surface and colored in white, black, and cyan. One monomer of *Mt*Fsr is represented as a cartoon and colored accordingly to the top panel. Clusters are numbered based on their position in the electron relay going from the FAD to the siroheme. The siroheme and FAD are represented in sticks and the [4Fe–4S]-clusters are shown as balls. Carbon, nitrogen, oxygen, sulfur, and iron atoms are colored respectively as purple (siroheme)/yellow (FAD), blue, red, yellow, and orange.

Here, we present the X-ray crystal structures of Fsr isolated from two *Methanococcales* as well as the EPR spectroscopy characterization of its metallocofactors, providing the first snapshots and molecular insights of this elegant machinery.

## Results

### Fsr conveys SO_3_^2-^ resistance in *M. thermolithotrophicus*

Fsr from *M. jannaschii* (*Mj*Fsr) has already been characterized by Johnson and Mhukopadhyay (4, 7, 19) and was firstly used in our study to obtain structural information. Since *Mj*Fsr had a crystallization defect (twinning, see materials and methods) we used a second model organism belonging to *Methanococcales*. Daniels *et al.* demonstrated that *Methanothermococcus thermolithotrophicus*, a marine thermophile isolated from geothermal heated sea sediments, could grow on 1 mM SO_3_^2-^ as a sole sulfur source, but growth was inhibited at higher concentrations (26, 27). Participation of Fsr in this process has not been examined before and was firstly investigated. After some adaptation, *M. thermolithotrophicus* could still grow on SO_3_^2-^ concentrations up to 40 mM (Fig. S2A). Accordingly, we measured a sulfite reductase activity 7.4 times higher in the cell extract of SO_3_^2-^ exposed cells than in the cell extract from a HS^-^-grown culture (2.95 compared to 0.4 μmol of reduced methyl-viologen oxidized.min^-1^.mg^-1^ of total protein). The soluble protein fraction of *M. jannaschii* and *M. thermolithotrophicus* grown on either HS^-^ or SO_3_^2-^ was compared on clear native PAGE (Fig. S2B). Upon SO_3_^2-^ exposure, a band of ≈ 300 kDa is produced with cellular levels comparable to those of the Methyl-Coenzyme M reductase (MCR), the main catabolic enzyme of the methanogenesis pathway (4). The same band was observed in the first report of Johnson and Mukhopadhyay (4) and was suspected to be Fsr. Since the *fsr* sequence was in the uncovered region of the shotgun genome of *M. thermolithotrophicus* (assembly number ASM37696v1, Bioproject: PRJNA182394), we sequenced the genome of the strain DSM 2095. The closed circular genomic sequence was obtained (see supporting information) and contained the whole *fsr* gene. The gene codes for a 69-kDa protein (abbreviated as *Mt*Fsr) that shares 80.4 % identity with *Mj*Fsr and belongs to Group I.

Fsr from both organisms were natively purified under anaerobic atmosphere and yellow light. SDS-PAGE profiles and sulfite reductase activity assays were used to track the enzyme during the purification (see Fig. S2C and materials and methods). *Mt*Fsr exhibits the typical absorbance of [Fe–S]-clusters and siroheme–[4Fe–4S] containing proteins as previously shown for *Mj*Fsr (Fig. S3A (4, 28)). Based on the clear native PAGE and gel filtration profiles, the purified *Mt*Fsr is organized as a tetramer (Fig. S2B and S3B). Accordingly, the gel filtration of *Mj*Fsr performed by Johnson and Mukhopadhyay suggested a penta- or tetrameric arrangement and we concluded that Fsr should form a homotetramer in solution.

### Inserted ferredoxin domains stabilize a tetrameric organization

The purified proteins were immediately used for crystallization and yielded brown crystals after several days. The first Fsr structure, obtained from *Mj*Fsr, was solved *ab initio* by a single-wavelength anomalous dispersion experiment measured at the Fe K-edge. *Mt*Fsr was solved by molecular replacement, using *Mj*Fsr as a template. The crystal structures of *Mj*Fsr and *Mt*Fsr were respectively refined to 2.30 Å (Fig. S4A) and 1.55 Å (Fig. 1 and Table S1). Both structures superpose well (Fig. S4B) and have an overall butterfly shape. Because *Mj*Fsr has a pseudo-merohedral twinning and a lower resolution compared to *Mt*Fsr, the latter was used for in depth structural analysis.

The monomeric architecture is organized as follows: a N-terminal ferredoxin domain containing two [4Fe–4S]-clusters (*Mt*Fsr residue 1-57), connected to the F_420_H_2_–oxidase domain that harbors the flavin and one [4Fe–4S]-cluster (*Mt*Fsr residue 58-336), linked to the C-terminal sulfite reductase domain (*Mt*Fsr residue 339-484, 546-618), which binds the siroheme–[4Fe–4S] (Fig. 1). The sulfite reductase domain has an inserted ferredoxin domain that contains two [4Fe–4S]-clusters (*Mt*Fsr residue 485-545). The tetramer is organized as a dimer of dimer. The basic monomer-monomer interface of 2,902-Å^2^ for *Mt*Fsr and 2,971-Å^2^ for *Mj*Fsr (Fig. S5) is established by the sulfite reductase domain and the two additional ferredoxin domains. The C-terminal part of the sulfite reductase domain (562-618 in *Mt*Fsr, 562-620 in *Mj*Fsr), the second ferredoxin domain and the loop 171-189 of the F_420_H_2_-oxidase domain generate the dimer-dimer interface, totaling an area of 3,055-Å^2^ for *Mt*Fsr and 3,037-Å^2^ for *Mj*Fsr (Fig. S5). Most of these contacts involve salt bridges. In *Mj*Fsr, the tetrameric structure is supported by two divalent cations, modelled as calcium ions that are each coordinated by a conserved aspartate from the opposite monomers (Asp511 and waters, Fig. S4A).

Because of its non-crystallographic symmetry and the four tetramers in the asymmetric unit, a total number of 96 [4Fe–4S]-clusters were modelled in the crystal structure of *Mt*Fsr, which, to our knowledge, is the highest number of clusters seen in an asymmetric unit so far (Fig. S6A). As shown by the B-factor profile, the sulfite reductase domain is very stable whereas the peripheral F_420_H_2_-oxidase domain shows a more dynamic behavior (Fig. S6B-C), especially in its lid region (204-253 for *Mt*Fsr and *Mj*Fsr). In *Mt*Fsr, one chain (chain N) seems to have two different conformations in the lid region, and both were tentatively modelled.

### Organization of the F_420_H_2_-oxidase domain

The F_420_H_2_-oxidase domain of Fsr is almost identical between *Mj*Fsr and *Mt*Fsr (rmsd=0.33 Å for 277-Cα aligned) and it superposes well to FrhB from *Methanothermobacter marburgensis* (PDB code: 4OMF (23), with a rmsd=0.92 Å for 179-Cα aligned) and to FrhB from *Methanosarcina barkeri* (PDB code: 6QGR (25), with a rmsd=0.98 Å for 179-Cα aligned) (Fig. 2A). The overall fold is perfectly conserved between the F_420_H_2_-oxidase domain of Fsr and FrhB, except for the helix α1 of FrhB that became a loop in Fsr (Fig. 2A).

**Figure 2.**
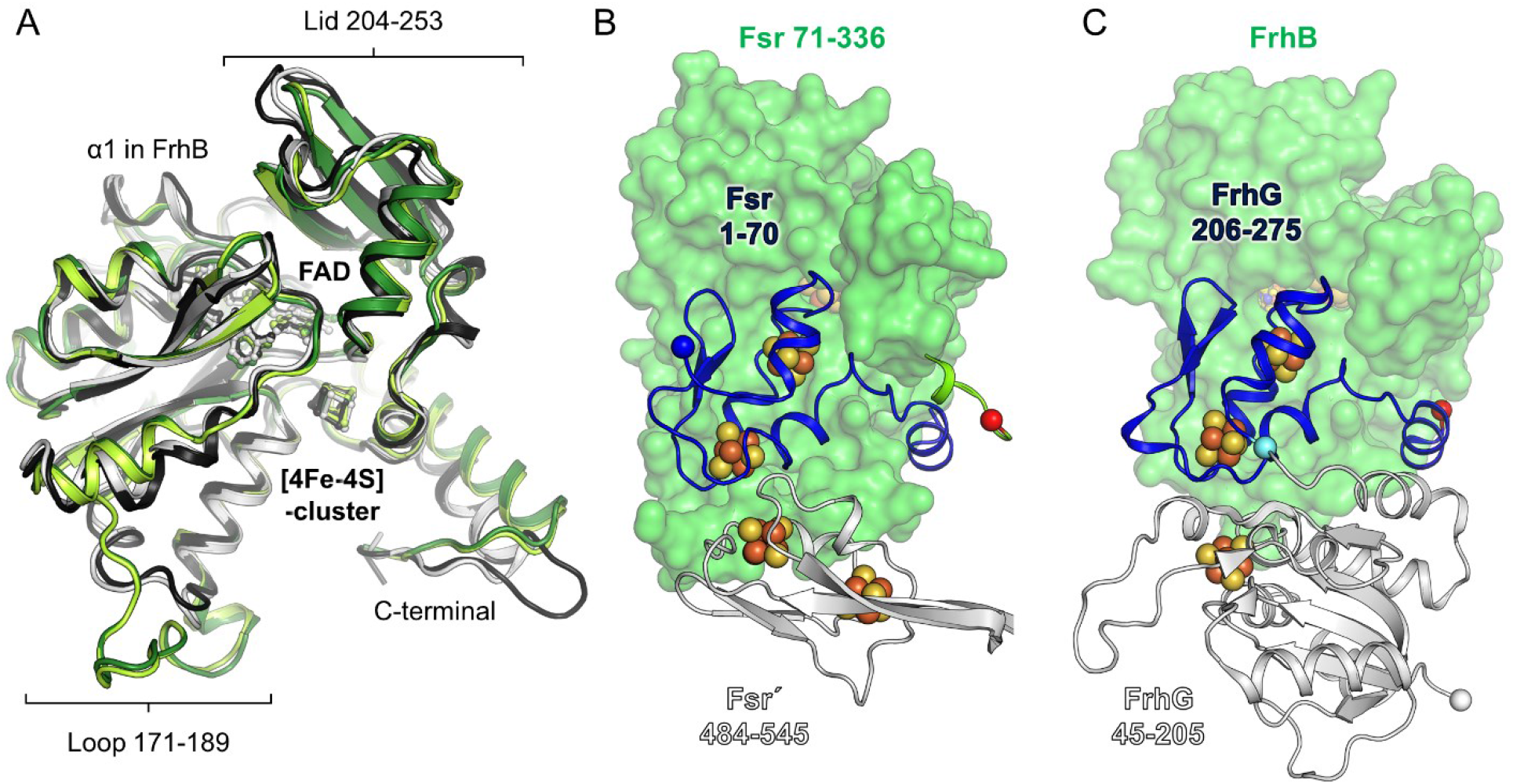
Comparison of the F_420_H_2_-oxidase domain between Fsr and Frh. (A) Superposition of the F_420_H_2_-oxidase domain in Fsr (*Mj*Fsr in dark green, *Mt*Fsr in light green) with FrhB from *M. barkeri* (black, PDB: 6QGR) and FrhB from *M. marburgensis* (white, PDB: 4OMF). The extended loops 171-189 in *Mj*Fsr and *Mt*Fsr are highlighted, as well as the lid, static in the Frh structures, but more flexible in Fsr (Fig. S6). (B) Representation of *Mt*Fsr F_420_H_2_-oxidase domain (green surface) and its N-terminal ferredoxin domain (blue cartoon residues 1-70). N-terminus of Fsr and C-terminus from the F_420_H_2_-oxidase domain are highlighted by blue and red spheres, respectively. The inserted ferredoxin domain, provided by the opposing monomer (Fsr’), is shown in white cartoon representation. (C) Arrangement of FrhB (green surface) with FrhG (cartoon) from *M. marburgensis* (PDB 4OMF). The N-terminal part (45-205) of FrhG is colored in white and its C-terminal part (206-275), structurally equivalent to the N-terminal ferredoxin domain of Fsr, is colored in blue. The cyan ball highlights the connection between both FrhG parts.

The active site contains a flavin adenine dinucleotide (FAD), which is bound in a similar fashion in Fsr and FrhB (Fig. S7A-B). Additional electron density was not observed despite the co-crystallization with reduced F_420_. The F_420_ coenzyme is supposed to bind in a positively charged cleft that complements the charges of the acidic gamma-carboxy groups (Fig. S7E) (23, 29). While the lid region (204-253) is fixed in the FrhB structures, possibly because of its interaction in an inter-subunit contact, it is more flexible in Fsr (Fig. S7B-C).

In Fsr, the N-terminal ferredoxin domain serves as an electron connector. In the Frh system, however, the small subunit of the [NiFe]-hydrogenase (FrhG) is sitting on the surface of FrhB and transfers the electrons. Surprisingly, despite such differences, FrhG and the N-terminal ferredoxin domain of Fsr are located at the same position on the F_420_-oxido/reductase domain (Fig. 2B-C). The extended loop 171-189 in *Mt*Fsr and *Mj*Fsr serves as a platform to specifically bind both ferredoxin domains. The Glu180, part on this loop, binds the [4Fe–4S]-cluster 3 (monodentate, 2.22 Å, Fig. 3 and Fig. S8-9). This is a distinct feature compared to the Frh system, where FrhG binds its three [4Fe–4S]-cluster without FrhB being involved.

**Figure 3.**
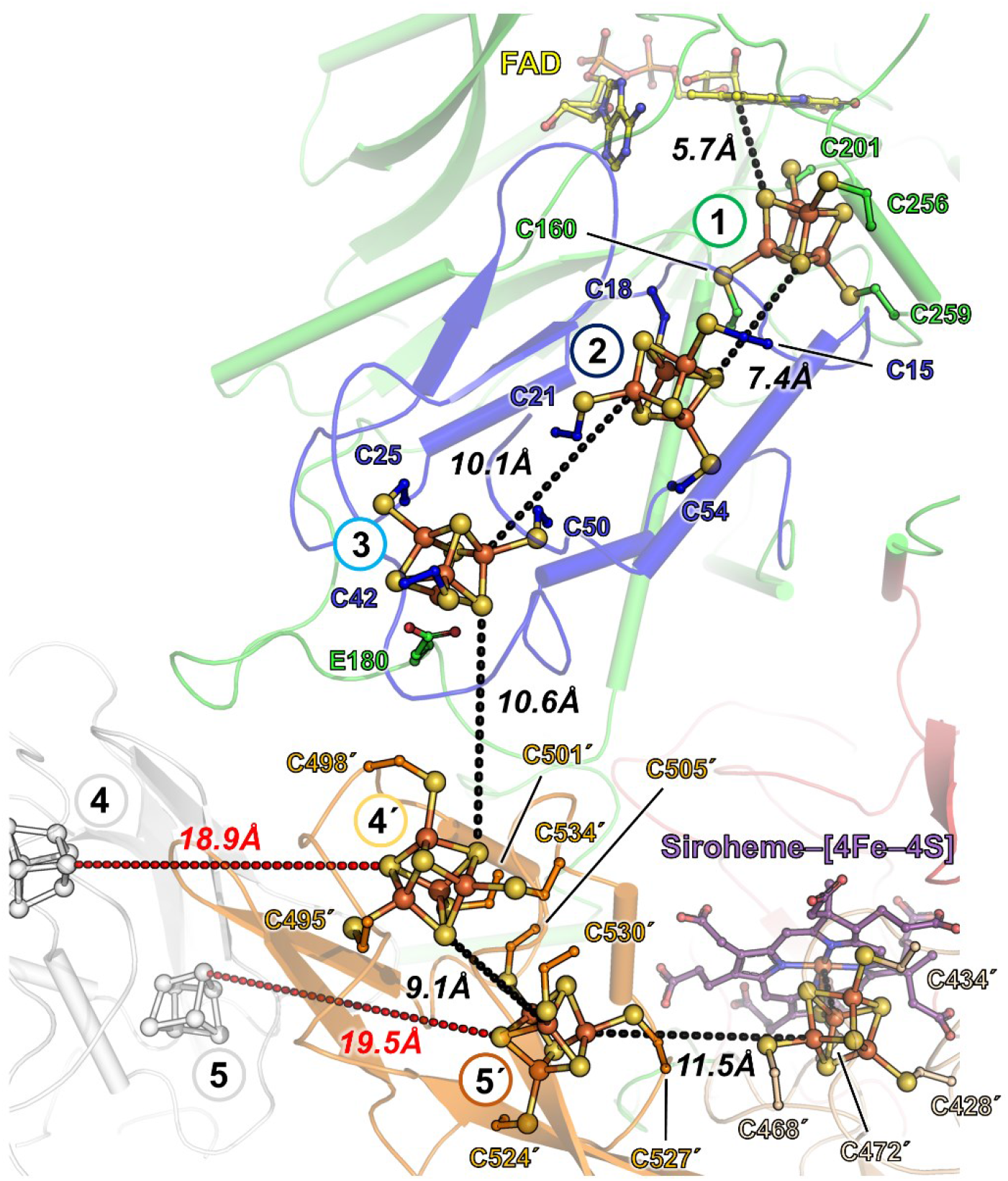
Electron transfer relay of *Mt*Fsr. *Mt*Fsr, shown as a cartoon has the same color code and numbering of its [4Fe–4S]-clusters (balls and sticks) as in Fig. 1. Edge to edge distances connecting the clusters are shown as dashes. The distances to the adjacent [4Fe–4S]-clusters of the opposite dimer are shown in red. The opposing monomer residues and clusters are labelled with prime symbol. The primes correspond to the second monomer forming the dimer. The residues binding the clusters are shown as balls and sticks. Carbon atoms are colored by their domain affiliation. Nitrogen, oxygen, sulfur, and iron atoms are colored in blue, red, yellow, and orange, respectively. Siroheme and FAD are shown as sticks with purple and yellow carbon atoms, respectively.

### Both active centers are electronically connected via a five [4Fe–4S]-cluster relay

The distance between the isoalloxazine ring from the FAD to the closest siroheme–[4Fe–4S] is circa 40-Å. Electrons delivered by reduced F_420_ must therefore travel through an electron-transfer relay to reduce SO_3_^2-^. As illustrated in Fig. 3, this path is constituted of five [4Fe–4S]-clusters electronically connected by edge-to-edge short distances (< 11.5 Å). The dimerization is critical to establish the overall electron path since half of the relay is provided by the second protomer. An intra-electron transfer between both dimers is unlikely due to the long distance that separates the nearest-by clusters (i.e., 18.9 and 19.5 Å).

Electrons on the isoalloxazine ring can be directly transferred to the [4Fe–4S]-cluster 1, located in the F_420_H_2_-oxidase domain. From here, they will be passed on to cluster 2 and 3 from the N-terminal ferredoxin domain, followed by cluster 4’ and 5’ from the inserted ferredoxin domain of the inter-dimer and finally arrive at the siroheme–[4Fe–4S]. Sequence analyses already predicted a total of four [4Fe–4S]-clusters and the one coupled to the siroheme (4). The extra cluster ([4Fe–4S]-cluster 1), discovered in both Fsr structures, shows a non-canonical binding sequence (PCX_40_CX_54_CX_2_C, Fig. S9). More importantly, the predicted four clusters have different ligands compared to what was previously proposed (Fig. S9). Each [4Fe–4S]-cluster has a different protein environment: Cluster 1 is surrounded by basic residues, cluster 2 and 5 have a hydrophobic shell, cluster 4 and 6 are in a more polar environment and cluster 3 has a glutamate ligand. These differences could reflect the need to establish a “redox potential ladder” to allow a smooth one-way transfer of the electrons. To investigate the electron transfer path, electrochemical experiments followed by Electron paramagnetic resonance (EPR) spectroscopy were performed.

### Redox properties of the seven metallocofactors

EPR spectroscopy at 10 K (Fig. S10) revealed that in as isolated *Mt*Fsr high spin (*S*=5/2) and low spin (*S*=1/2) signals typical for the siroheme in sulfite reductases (30, 31) were absent, neglecting the sharp axial *S*=5/2 EPR signal around *g*=6, which quantified by double integration of its simulation spectrum (*g*=6.22, 5.92 and 1.98) is at most 3 % of *Mt*Fsr. Apparently, upon purification under strictly anaerobic conditions, the siroheme remains in its ferrous state. After methylene blue oxidation or upon dye-mediated redox titration with *E*_m,7.5_ = −104 mV versus H_2_/H^+^ (Normal hydrogen electrode, NHE) an intense rhombic *S*=5/2 EPR signal with *g*=6.7 and 5.1 appeared (Fig. 4A-B). The spectrum could be simulated with three components: a main species with *g*=6.70 and 5.10 (78 %), a less abundant species (19 %) with almost identical *g*-values (6.80 and 5.08) but a narrower linewidth and the sharp axial *g*=6 species already occurring in the as isolated *Mt*Fsr. For both rhombic components *g*=1.95 was taken as third *g*-value, as the experimental spectrum contained a weak [3Fe-4S]^1+^ signal from limited [4Fe–4S]^2+^ breakdown upon oxidation. In sulfite reductase and other hemoproteins the occurrence of multiple high spin species is not unusual (32). Addition of sulfite to methylene blue oxidized *Mt*Fsr led to disappearance of the siroheme ferric high spin signals and formation of a weak low spin EPR signal of which only the highest *g*-value (2.8) was detectable, in agreement with the known reactivity of sulfite reductases (33).

**Figure 4.**
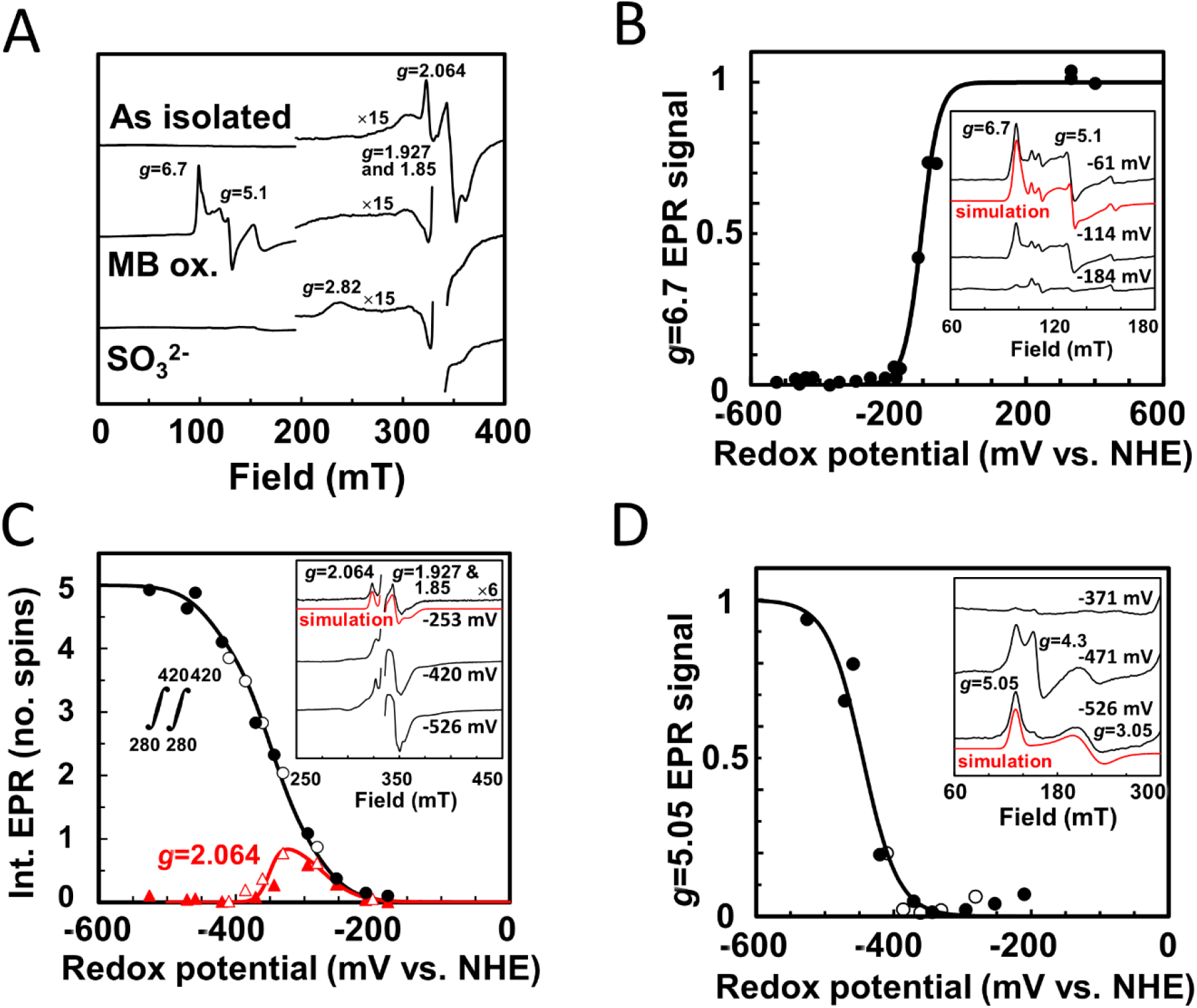
Determination of the redox potential of the metallocofactors in *Mt*Fsr via EPR spectroscopy. (A) EPR spectra of as isolated, methylene blue (MB) oxidized and, consecutively, sodium sulfite (10 mM) treated *Mt*Fsr. (B-D) Dye-mediated redox titrations of indicated EPR signals (or double integral in (C)). Representative spectra at three selected potentials are shown in the insets, including *g*-values and simulations (see text). EPR spectra for all samples are in Fig. S10. Nernst fits for n=1 with *E*m= −104 mV (B), −275, three times −350 and −435 mV (C) and −445 mV (D) are shown. The fit for *g*=2.064 used n=1 (in red) for −275 mV and n=2 for −350 mV (in black). EPR conditions: temperature, 10 K; modulation frequency, 100 kHz; modulation amplitude, 1.0 mT; microwave frequency 9.353 GHz; microwave power 20 mW except in (C), 0.2 mW.

In an enzyme approaching the complexity of the complex I it is not feasible to determine all individual redox potentials of its five regular [4Fe–4S]^1+/2+^ cubanes and the cubane bridged to siroheme. First, based on the distances in Fsr, extensive magnetic coupling (refer to (34)) between neighboring cubanes is anticipated, which blurs EPR spectroscopic features of individual [4Fe–4S]^1+^ clusters. Second, the coupling between the ferrous siroheme and its cysteine-bridged reduced cubane leads to complex mixtures of sharp *g*=1.94, broader *g*=2.29 and very anisotropic *S*=3/2 mimicking signals (35). Third, we had to master the administration of reducing equivalents to Fsr at known solution potential, avoiding sodium dithionite with its degradation product sulfite. With the sodium borohydride reduced natural electron donor F_420_ (36) electrons could conveniently be introduced, while following the solution potential with mediators. One [4Fe–4S]^1+/2+^ cubane with *g*-values simulated with 2.064, 1.927 and 1.85 was already reduced at a relatively high potential and is also detected in as isolated Fsr (Fig. 4A and C). By following the amplitude of the second derivative of the experimental EPR spectrum at *g*=2.064 *E*_m,7.5_ = −275 mV NHE was estimated from fitting to the Nernst equation with n=1 (Fig. 4C). The signal “disappeared” upon further reduction with *E*_m,7.5_ = −350 mV NHE in a manner which indicated cooperativity (n≥2). As super-reduction to [4Fe–4S]^0^ is unlikely (*E*_m_ = −790 mV (37)), we interpret this phenomenon as reduction of two neighboring clusters of the *g*=2.064 cluster. This cluster thus is number 2, 3 or 4’ (the siroheme cubane typically has a very low potential (30)). In the absence of other well-defined EPR features below −350 mV we double integrated the EPR spectra. Based on iron-content divided by 24 (siroheme does not release Fe-ions in acid) we quantified the spin concentration at the lowest attainable potential (−526 mV) to be 4.5 ± 0.5 spins, which most likely correspond to the five regular clusters. A fit for the spin integral as function of the redox potential had to include the experimental *E*_m,7.5_ = −275 mV NHE and *E*_m,7.5_ = −350 mV NHE for both neighboring clusters.

Without overfitting we could satisfactorily reproduce the data for five redox transitions at three midpoint potentials: one at *E*_m,7.5_ = −275 mV NHE (experimental), one at a low potential to represent the lowest potential region (*E*_m,7.5_ = −435 mV NHE) and three times *E*_m,7.5_ = −350 mV NHE for the other three clusters (which includes the two clusters leading to broadening of the *g*=2.064 signal).

In the low field region of the EPR spectrum, a species with unusual *g*-values was detected (simulated *g*-values 5.05, 3.05 and 1.96) at very low potential (Fig. 4D). It was accompanied in some samples by an isotropic *g*=4.3 signal. But since the integrated intensity was maximally 5% of the *g*=5.05 species and non-Nernstian behavior was seen; it was not considered physiologically relevant. From the work of Christner et al. (35) there is no doubt that the *g*=5.05 species is not from a *S*=3/2 system but from transitions of the siroheme-Fe^2+^ exchange-coupled to the [4Fe–4S]^1+^ cubane with J/D≈ −0.2 and E/D≈ 0.11 (Fig. S10C). In full agreement with findings on the *E. coli* assimilatory reductase (30) a very low potential (*E*_m,7.5_ = −445 mV NHE) was estimated.

### An elementary sulfite reductase

The C-terminal domain of Fsr represents the most simplified sulfite reductase crystallized so far. While sharing the common fold of sulfite reductases (Fig. S11, Fig. S12) (10, 15, 16), Fsr lacks the large N- and C-terminal extensions found in aSirs and dSirs, presumably used to fortify dimerization and maybe to interact with partners. Without extensions, Fsr is much more simplified and compact, possibly a thermophilic trait. Each Fsr protomer contains one functional siroheme center, in comparison to dSirs that harbor one functional and one structural siroheme center in each DsrAB heterodimer, or aSirs that have lost one siroheme–[4Fe–4S] site (Fig. 5 and S12).

**Figure 5.**
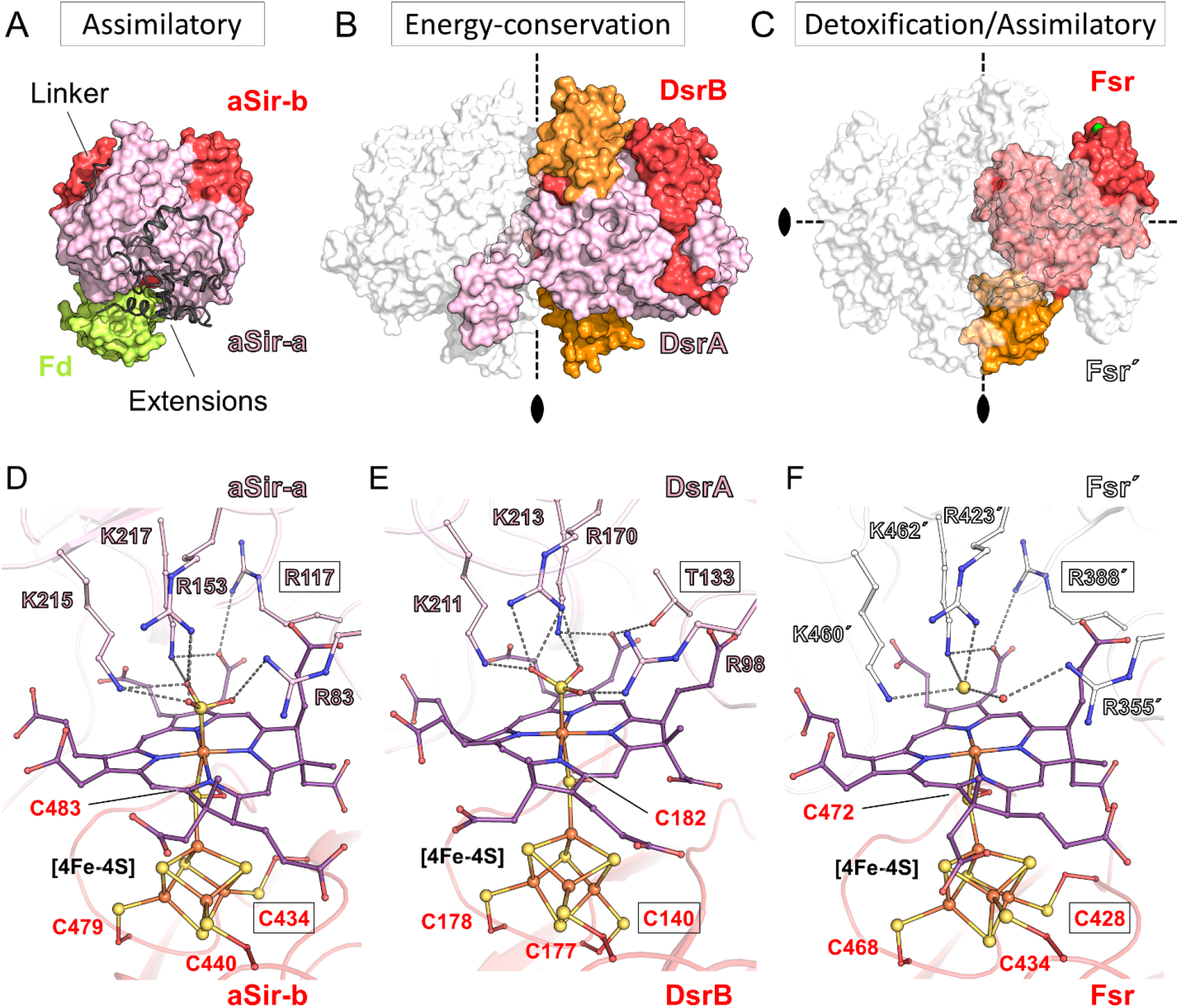
Overall structural comparison between aSir, dSir and Fsr. (A-C) All structures are represented in surface, dimeric partners shown in white transparent and residues from the opposing monomer are labelled with prime symbol. The inserted ferredoxin domains of DsrAB and *Mt*Fsr are colored in orange. (A) aSir from *Zea mays* with its [2Fe–2S]-ferredoxin colored in light green (PDB 5H92). (B) DsrAB from *A. fulgidus* (PDB 3MM5) and (C) *Mt*Fsr tetramer. For *Mt*Fsr, the green surface indicates the F_420_H_2_-oxidase position. (D-F) Active site of sulfite reductases. Close up of the active site and the functional siroheme surrounding in (D) *E. coli* aSir (PDB 1AOP), (E) dSir of *A. fulgidus* (PDB 3MM5), (F) *Mt*Fsr in which sulfide was tentatively modelled. Residues coordinating the [4Fe–4S]-cluster, the siroheme and the sulfur species are shown as balls and sticks, while sulfur and iron are depicted as spheres. Framed residues highlight the differences between the siroheme–[4Fe–4S] binding in aSirs and dSirs.

Despite being phylogenetically more distant from aSirs than to dSirs, Fsr superposes relatively well to the first and second half of aSirs (Fig. S11, S13-14). The position of the C-terminus of Fsr matches the beginning of the linker, which connects the two halve domains in aSir (Fig. S11-S14). This detail corroborates the theory that modern aSirs evolved through duplication and fusion events.

The inserted ferredoxin domain in Fsr is at the same position as the ferredoxin domain in DsrA or DsrB (Fig. S11-12, Fig. S15-16). There is a remarkable three-dimensional conservation of the electron connectors between the structures of Fsr, DsrA, DsrB and even the aSir from *Zea mays*, where the external [2Fe–2S] ferredoxin (PDB 5H92 (38)) sits on the core of the sulfite reductase (Fig. 5 and S12). Such a conserved position suggests a common origin. Alternatively, this conserved arrangement might be due to the restricted access to the cluster bound to the siroheme, and the selective pressure towards the most optimized distance for the electron transfer.

### Fsr has structural and catalytic traits of assimilatory sulfite reductases

While the sirohemes of DsrAB are partially surface exposed in order to interact with DsrC (Fig. S1, (15)), the sirohemes of Fsr are buried within the sulfite reductase domain but still accessible via a positively charged solvent channel (Fig. S17A). As in DsrAB, the two sirohemes within one Fsr dimer are in close proximity (9.4 Å, Fig. S17B (16)).

The binding of the siroheme in *Mj*Fsr and *Mt*Fsr is highly conserved (Fig. S18). It is mainly anchored by positively charged residues from one protomer while the dimeric partner binds the adjacent [4Fe–4S]-cluster, to establish the siroheme–[4Fe–4S] center, as reported for other sulfite reductases. Based on the observed electron density, we tentatively modelled a SO_3_^2-^ ion bound to the siroheme-iron (short distance of 2.3 Å, Fig. S19A) in *Mj*Fsr. In *Mt*Fsr, the axial ligand is a single atomic species observed in the 16 sites of the asymmetric unit, which is not covalently bound to the iron (distance of 2.9 Å, Fig. S19B). An anion HS^-^ was modelled in the electron density based on the pH of the crystallization solution (pH of 5.5). This species might be the result of co-crystallizing Fsr with reduced F_420_, which could have forced the complete reduction of bound SO_3_^2-^. Superposition analysis shows that the functional sirohemes are arranged in a highly similar manner, whereas the binding of the structural siroheme or sirohydrochlorin differ, highlighting the strong influence of the protein environment on the siroheme geometry (Fig. S19C).

Four positively charged residues (Arg423, Arg355, Lys460 and Lys462), perfectly conserved across sulfite reductases (Fig. 5, S13 and S15), bind the SO_3_^2-^ as well as two water molecules in *Mj*Fsr. This solvent network is suspected to allow a faster proton transfer via the Grotthuss mechanism, required for the reduction reaction as proposed in dSirs (18, 39). In *Mt*Fsr, the modelled sulfide ion is bound by the Arg423, Lys460, Lys462 and one water molecule stabilized by the Arg355. It is worth to note that Group II Fsr, found in the genome of ANME (except for *Ca*. *Methanoperedens nitroreducens*) and *Methanosarcinales*, should have a larger binding pocket. In Group II Fsr two of the SO_3_^2-^-binding arginine residues, found in Group I Fsr, are replaced by a lysine and glycine. This suggests a different substrate specificity for this functionally uncharacterized enzyme (5).

Structural comparison of the sulfite reductase active sites presented two critical positions: Firstly, a canonical threonine in DsrA (αThr136 in *D. vulgaris* and αThr133 in *A. fulgidus)* is absent in Fsr. Instead, it is replaced by an arginine (Arg388), which is a hallmark of the aSir family. Secondly, the [4Fe–4S]-cluster coupled to the siroheme is coordinated in the exact same way as aSirs and DsrA, with the canonical motif: CX_5_CX_n_CX_3_C. In comparison, the catalytically active [4Fe–4S]-cluster coupled siroheme of DsrB exhibits the canonical motif CX_n_CCX_3_C (Fig. 5 and Fig. S13-16). Based on these observations, Fsr must adopt the same catalytic path as aSirs. Therefore, it should perform a unidirectional and complete six electron reduction of SO_3_^2-^ to S^2-^ without the formation of intermediates or side products. Johnson and Mukhopadhyay previously reported that *Mj*Fsr does not reduce thiosulfate, an intermediate of the trithionate pathway and characteristic trait of the dSir family (4).

We also monitored the F_420_ reduction by *Mt*Fsr with S^2-^ as a substrate and observed no reaction even at 10 mM of S^2-^ concentration. Also, the addition of 10 mM S^2-^ to 1.4 mM of Na_2_SO_3_ had no impact on the F_420_H_2_-oxidation rate. The apparent *Km* value for SO_3_^2-^ was 15 ± 2 μM, and the corresponding app *V_max_* value was 27 ± 1 μmol of F_420_H_2_ oxidized.min^-1^.mg^-1^ of pure *Mt*Fsr (see Materials and methods). *Mt*Fsr was also able to use nitrite to oxidize F_420_H_2_ with an app*K*_m_ value for NO_2_^-^ of 2.5 ± 0.2 μM, and the corresponding app *V_max_* value was 27 ± 0.5 μmol of F_420_H_2_ oxidized.min^-1^.mg^-1^ of pure *Mt*Fsr. Additionally, *Mt*Fsr was shown to reduce selenite (SeO3^2-^) with a relative activity of 20.7 ± 7.5 % compared to SO_3_^2-^ (see Materials and methods). *Mt*Fsr did not accept thiosulfate as an electron acceptor, supporting that, indeed, Fsr is acting like an aSir.

## Discussion

Some methanogens show a remarkable tolerance to SO_3_^2-^, one of the sulfur reactive species that can cause oxidative damage to the methanogenic machinery. Besides the possibility that those methanogens can keep low intracellular SO_3_^2-^ concentrations through pumping mechanisms, the cytoplasmic Fsr type I is used as a first line of defense. The enzyme converts the poisonous SO_3_^2-^ to HS^-^, which can be further used for S-assimilation. Previous studies have shown that *M. jannaschii* is rapidly expressing a vast amount of Fsr during growth of SO_3_^2-^ (4) and our work showed the same response in *M. thermolithotrophicus*. Fsr corresponds to 5–10 % of cellular protein (calculated from Fsr’s activity in the cell extract and the specific activity of the purified enzyme), an amount similar to methyl-CoM reductase (see Fig. S2B). Fsr is fueled by the electron donor F_420_H_2_, which is highly abundant in the cytoplasm (22) and that can be quickly reduced via H_2_-oxidation through the Frh machinery.

In contrast to the bi-directional hydrogenase Frh that maintains an isopotential of *E*°′≈ −400mV (23), the different metallo-clusters of Fsr must establish a downhill redox-potential from the FAD to the siroheme–[4Fe–4S] (Fig. 6). Our electrochemical and spectroscopic studies indicate that electrons carried by F_420_H_2_ are immediately transferred to the siroheme–[4Fe–4S] (Fig. 4 and S10). The environment of the metallo-cofactors should assure the efficient electron transfer rather than serving as a transient storage and a cascade of redox potential from −380 mV (F_420_/F_420_H_2_ redox potential under certain physiological conditions (22)) to −116 mV (*E*′° of HSO_3_^-^/HS^-^) is expected. In contrast, while one cluster has indeed a measured redox potential of −275 mV and three others are at −350 mV, one of them exhibits a lower potential of −435 mV. The presence of such a low redox potential cluster has already been seen in the complex I and is not contradicting our proposed hypothesis regarding the electron flow. The particular nature and role of this cluster in the electron chain would be interesting to localize. Unfortunately, dissection by heterologous expression of domains is impossible, due to the intertwined nature of the electron transfer path and essential subunit interactions within the tetramer.

**Figure 6.**
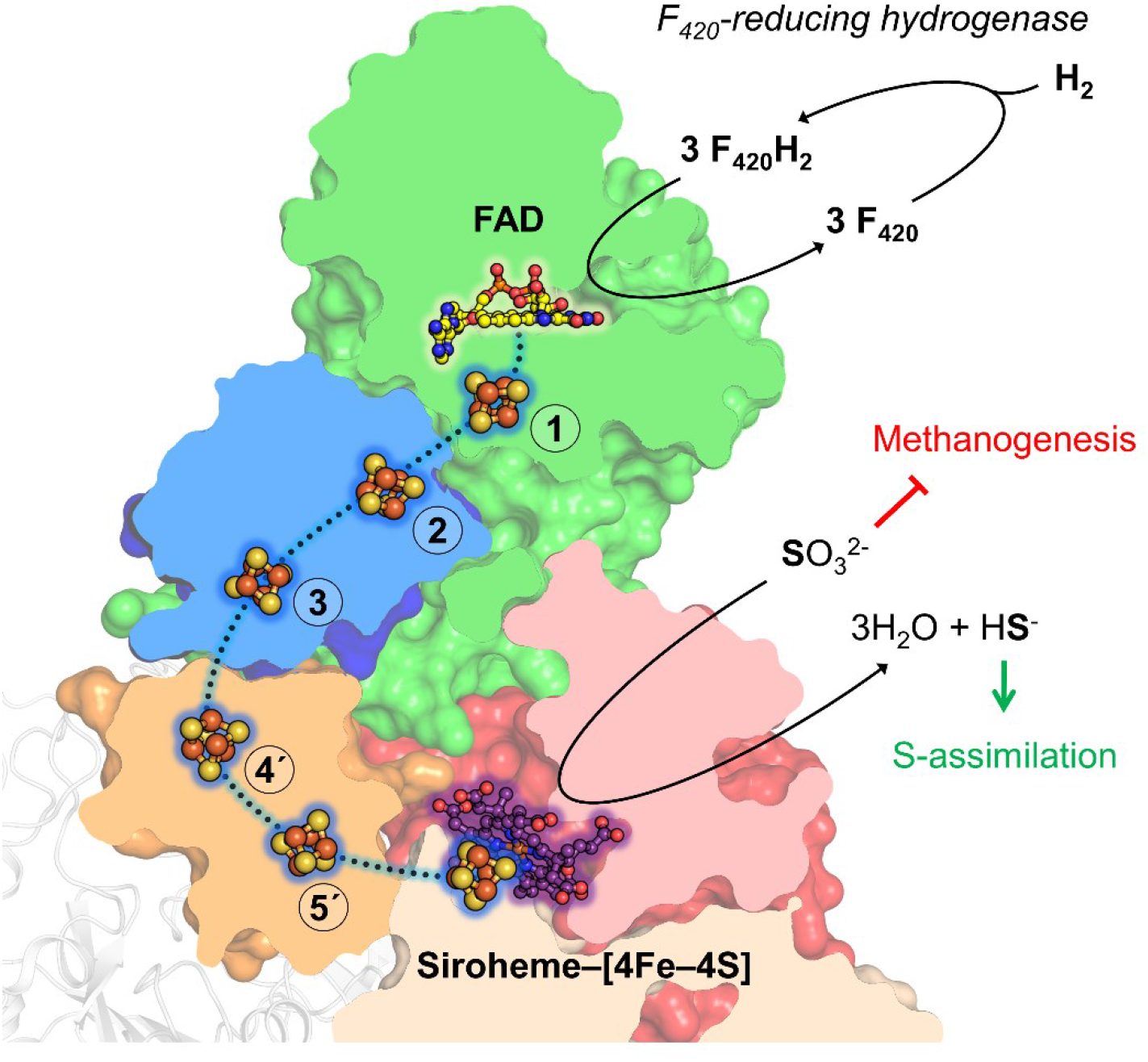
Overall view of the F_420_H_2_-based SO_3_^2-^-reduction reaction in hydrogenotrophic methanogens. Reduced F_420_ delivers two electrons to the FAD in the F_420_H_2_-oxidase domain of Fsr (green). Oxidized F_420_ will be reduced by Frh in hydrogenotrophic methanogens. The electrons harbored on the FAD will be transferred one by one on a five [4Fe–4S]-cluster relay, passing two ferredoxin domains (blue and orange) and will end up on the cluster coupled to the siroheme. The clusters labelled with a number and prime (4’, 5’ and the cluster coupled to the siroheme) are provided by the dimeric partner. The toxic SO_3_^2-^, inhibiting the methanogenesis, is electrostatically attracted to the siroheme–[4Fe–4S]. The successive reduction of SO_3_^2-^ leads to the formation of HS^-^ and H_2_O. HS^-^ will be either used for assimilatory purposes of the cell or released into the environment.

Once reduced, the siroheme–[4Fe–4S] could transfer the electrons to the sulfur species covalently bound on its Fe. dSirs physiologically perform a two-electron reduction to allow the transfer of the sulfur intermediate to DsrC. In contrast, aSirs and Fsr perform a three times two-electron reduction to release HS^-^. A positively charged environment surrounding the active site attracts SO_3_^2-^ and an organized water network would provide the protons required for its successive reduction (Fig. S17A). Despite a strikingly similar position of the residues involved in the substrate binding, aSirs/Fsr and dSirs are reacting differently. With the possibility to genetically modify *M. maripaludis* or *M. jannaschii* it would be worthwhile to exchange the residues that confer aSir traits at the active site (Arg388, Cys428) with dSir ones and observe the repercussion on the phenotype (7, 8).

Fsr appears not to be entirely substrate specific, as it is able to reduce selenite and nitrite. Nitrite and sulfite reductases share a similar reaction mechanism (12) and the kinetic parameters measured for nitrite suggest that Fsr might also be used for the conversion of this compound to ammonium. Interestingly, there is one report showing that *M. thermolithotrophicus* can grow on nitrate as a sole nitrogen source (40). Since the reduction of nitrate results in nitrite formation, *M. thermolithotrophicus* needs an efficient system to convert the toxic compound. Fsr would be a promising candidate since no nitrite reductase can be found in the genome. However, since hydrothermal fluids are usually depleted in both nitrate and nitrite (1, 41), we cannot exclude that the low app*K*_m_ value obtained for nitrite is due to substrate ambiguity rather than being physiologically important.

Based on the sequence and structural similarity to enzymes from different superfamilies it has been proposed that modern sulfite reductases originated from a primordial Sir/Nir functioning as a homodimer (9). A snapshot of this progenitor can be derived from the Fsr structure because of the very simplified organization of its sulfite reductase domain (Fig. S20). In this model, the homodimer would complement itself: one monomer would harbor the siroheme–[4Fe–4S], whereas the second monomer would contain the substrate/siroheme binding site, and reciprocally. The different steps that led to the evolution from this progenitor to modern Fsr can be hypothesized based on its modular organization. Following a simplistic model, a ferredoxin containing the 2 x [4Fe–4S] cluster would have been inserted in the elementary sulfite reductase module. Then, a F_420_H_2_-oxidase containing a ferredoxin domain (Fqo/FpoF like) would have fused to the N-terminus of the sulfite reductase domain containing the inserted ferredoxin (Fig. S18). Some members of the Sir superfamily might have derived from one of these steps (9), the similarities between the quaternary organization of Fsr and DsrAB and the active site of Fsr and aSir are particularly striking.

The evolution of Fsr is still a matter of debate and needs to be thoroughly studied because its discovery reinforced the question whether sulfate-respiration or methanogenesis was the primeval means of energy conservation during the evolution of early Archaea (42, 43). Both metabolisms are related to each other, and it is possible that they co-existed or even co-exist (44). Methanogens might have lost the genes required to perform the complete sulfate dissimilation over time but kept the sulfite reductase to adapt to environments where SO_3_^2-^ fluctuations do occur.

There is accumulating evidence that methanogenesis and sulfate reduction were and still are intertwined pathways in ancient archaea (5, 9). While so far, no archaea have been experimentally proven to perform both catabolic processes, *M. thermolithotrophicus* has been shown to grow on sulfate as a sole sulfur source (26). The final step of this assimilatory pathway would lead to the accumulation of SO_3_^2-^ as an intermediate and Fsr is expected to orchestrate this last reaction. Even though further studies need to investigate if this methanogen can express other enzymes of the sulfate reduction pathway, the structural elucidation of Fsr provides the first snapshot of a sulfate reduction-associated enzyme in a methanogen.

## Material and Methods

### Methanogenic archaea strains and cultivation medium

*M. jannaschii* (DSM 2661) and *M. thermolithotrophicus* (DSM 2095) cells were obtained from the Leibniz Institute DSMZ-German Collection of Microorganisms and Cell Cultures (Braunschweig, Germany) and cultivated in a previously described minimal medium with some modifications (45). Refer to supporting information to find the complete composition of the medium used in this study.

### Anaerobic growth of *Methanococcales*

For all studied archaea, cell growth was measured spectrophotometrically by measuring the optical density at 600 nm (OD_600nm_). To control the purity of the culture, samples were taken and analyzed via light microscopy. Both methanogens were cultivated at 65 °C with 1 x 10^5^ Pa of H_2_/CO_2_ in the gas phase. *M. jannaschii* was cultivated in flasks and *M. thermolithotrophicus* was cultivated in flasks or fermenter. Refer to supporting information to find the complete cultivation protocol used in this study.

### Purification of Fsr

All steps were performed under the strict exclusion of oxygen and daylight. Protein purifications were carried out in a Coy tent with an N_2_ and H_2_ atmosphere (97:3) at 20 °C under yellow light. For both Fsr, three to five chromatography steps were used with some variations. Refer to supporting information to find the complete purification protocols used. Fsr purification was further followed via activity assays and based on absorbance peaks at the wavelengths of 280, 420 and 595 nm. Each elution profile was systematically controlled by SDS-PAGE to select the purest fractions.

### Protein crystallization

The purified enzymes were kept in 25 mM Tris/HCl pH 7.6, 10 % v/v glycerol and 2 mM dithiothreitol. Fresh, unfrozen samples were immediately used for crystallization. Crystals were obtained anaerobically (N_2_/H_2_, 97:3) by initial screening at 20 °C using the sitting drop method on 96-Well MRC 2-drop crystallization plates in polystyrene (SWISSCI) containing 90 μl of crystallization solution in the reservoir.

### Crystallization of *Mj*Fsr

*Mj*Fsr (0.5 μl) at a concentration of 6.1 mg.ml^-1^ was mixed with 0.5 μl reservoir solution. Black, long plate-shaped crystals appeared after a few days in the following crystallization condition: 45 % v/v 2-methyl-2,4-pentanediol, 100 mM Bis-Tris pH 5.5 and 200 mM calcium chloride.

### Crystallization of *Mt*Fsr

*Mt*Fsr at a concentration of 11 mg.ml^-1^ was co-crystallized with FAD (0.5 mM final concentration) and F_420_H_2_ (15.5 μM final concentration). The protein sample (0.6 μl) was mixed with 0.6 μl reservoir solution. Thick square-shaped brown crystals appeared after few days. The reservoir solution contained 200 mM lithium sulfate, 100 mM Bis-Tris, pH 5.5 and 25 % w/v polyethylene glycol 3350.

### X-ray crystallography and structural analysis

Crystal handling was done inside the Coy tent under anaerobic atmosphere (N_2_/H_2_, 97:3). *Mj*Fsr crystals were directly plunged in liquid nitrogen whereas *Mt*Fsr crystals were soaked in their crystallization solution supplemented with 20 % v/v ethylene glycol as a cryo-protectant prior to being frozen in liquid nitrogen. Crystals were tested and collected at 100 K at the Synchrotron Source optimisée de lumière d’énergie intermédiaire du LURE (SOLEIL), PROXIMA-1 beamline; the Swiss Light Source, X06DA – PXIII and at PETRA-III, P11.

### *Mj*Fsr

After an X-ray fluorescence spectrum on the Fe-K edge, datasets were collected at 1.74013-Å to perform the single-wavelength anomalous dispersion experiment. Native datasets were collected at a wavelength of 0.97857-Å on the same crystal. Data were processed and scaled with autoproc (46). Phasing (obtained maximum CFOM for the substructure determination was 69), density modification and automatic building was performed with CRANK-2 (47). The asymmetric unit of *Mj*Fsr contains two half homotetramers. The model was then manually built with COOT and further refined with PHENIX (48, 49). X-ray crystallographic data were twinned, and the refinement was performed by applying the following twin law - k, -h, -l. During the refinement translational-liberation screw was applied.

### *Mt*Fsr

Data were processed and scaled with *autoPROC*. The structure was solved by molecular replacement with phaser from Phenix, using *Mj*Fsr as a template (49). The asymmetric unit of *Mt*Fsr contains four homotetramers. This crystalline form presents a significant translational non-crystallographic symmetry (14 %). The model was then manually rebuilt with COOT and further refined with *PHENIX*. During the refinement, non-crystallographic symmetry and translational-liberation screw were applied. In the last refinement cycles, hydrogens were added in riding position. Hydrogens were omitted from the final deposited model.

Both models were validated through the MolProbity server (http://molprobity.biochem.duke.edu) (50). Data collection and refinement statistics, as well as PDB identification codes for the deposited models and structure factors are listed in Table S1. Figures were generated with PyMOL (Schrödinger, LLC). Structural comparison was performed with the dissimilatory sulfite reductases from: *Desulfovibrio vulgaris* (2V4J), *Archaeoglobus fulgidus* (3MM5) and with the assimilatory sulfite reductase from *Escherichia coli* (1AOP) and *Zea mays* (5H92).

### Enzymatic assays

Enzymatic Fsr measurements were performed in 200 mM KH_2_PO_4_ buffer pH 7.0 under strict exclusion of hydrogen and oxygen. F_420_ was reduced by Frh as stated in the supporting information. The oxidation of the reduced electron donor F_420_ was followed spectrophotometrically at 420 nm. For F_420_H_2_, a molecular extinction coefficient of 33.82 mM^-1^·cm^-1^ at 420 nm was experimentally determined for the above-mentioned conditions.

The assays for the specific enzyme activity were performed at 65 °C in a 1 ml quartz cuvette closed with a butyl rubber stopper. The gas phase of the cuvette was exchanged several times with N_2_. To monitor the reduction of SO_3_^2-^, 1.4 mM Na_2_SO_3_ and 47.3 μM F_420_H_2_ were added to the KH_2_PO_4_ buffer. Once the spectrophotometer (Agilent Cary 60 UV-Vis) displayed a stable signal, the reaction was started by the addition of 0.19 μg *Mt*Fsr. To investigate whether *Mt*Fsr can use substrates other than SO_3_^2-^, we provided 1.4 mM of disodium thiosulfate (S_2_O_3_^2-^), 1.4 mM sodium nitrite (NO_2_^-^) or 1.4 mM disodium selenite (SeO_3_^2-^). We further tested if *Mt*Fsr can function in the reverse way by providing 1.4 mM disodium sulfide (S^2-^) as an electron donor and 47.3 μM of oxidized F_420_. All experiments were performed in triplicates.

The app*K*_m_ and app *V_max_* of *Mt*Fsr for SO_3_^2-^ and NO_2_^-^ were determined at 50 °C under an anaerobic atmosphere (100% N_2_). The assays were performed in 96-deep well plates and spectrophotometrically monitored (FLUOstar Omega multi-mode Microplate Reader). To determine the app*K*_m_ and app *V_max_* of *Mt*Fsr, 0 - 500 μM Na_2_SO_3_ or NaNO_2_ and 50 μM F_420_H_2_ were added to the 200 mM KH_2_PO_4_ buffer pH 7.0 and the reaction was started by the addition of 3.8 ng *Mt*Fsr. All experiments were performed in triplicates.

### EPR spectroscopy

The midpoint potentials of the [4Fe–4S]-centers and the siroheme of *Mt*Fsr were determined from EPR signal intensities and EPR integrals of the various redox states. All titrations were performed in a Coy tent (N_2_/H_2_, 97:3), at 25 °C in the dark. A volume of 3.32 or 3 ml for the reductive or oxidative titrations with F_420_H_2_ or potassium ferricyanide at an initial *Mt*Fsr concentration of 4.07 or 2.7 mg/ml (in 100 mM Tris/HCl, pH 7.6), respectively, was stirred under anaerobic conditions. The solution potential was measured with an InLab ARGENTHAL (Mettler, Germany) microelectrode (Ag/AgCl, +207 mV versus H_2_/H^+^ with in-built platinum counter electrode) in the presence of the respective mediator mix. *Mt*Fsr was pre-incubated for 30 minutes before each titration with the mediator mix and assay buffer. The amount of *Mt*Fsr available and the necessary protein concentration to obtain a satisfying signal-to-noise for the EPR spectra precluded multiple titrations. Thus, values reported were from a single redox titration for the siroheme and from two redox titrations for the Fe/S signals.

The mediator mix for the reductive titration contained methylene blue, resorufin, indigo carmine, 2-hydroxy-1,4-naphthoquinone (50 μM), sodium anthraquinone-2-sulfonate, phenosafranin, safranin T, neutral red, benzyl- and methylviologen (all at a final concentration of 25 μM, except 2-hydroxy-1,4-naphthoquinone). For the oxidative titration the mediator mix contained methylene blue, resorufin, indigo carmine, 2-hydroxy-1,4-naphthoquinone (all at a final concentration of 20 μM). After adjustment of the potential by microliter additions of F_420_H_2_ or potassium ferricyanide and 3 minutes equilibration, EPR samples were taken. For this, 300 μl of the mix were withdrawn, removed from the anaerobic glove box in EPR tubes after attachment of a 5 cm piece of ID 3 mm x OD 7 mm natural rubber tubing sealed with a 5 mm OD acrylic glass stick at the other end. The samples were stored in liquid nitrogen until EPR spectra were recorded.

*Mt*Fsr as isolated was already in a partially reduced state. To obtain the completely oxidized form 675 μl Fsr at 20 mg/ml were incubated for 30 minutes with 2 mM methylene blue. The sample was then passed through a Sephadex G-25M column (previously equilibrated with 100 mM Tris/HCl pH 7.6) to remove the methylene blue. This methylene blue treated Fsr (1.28 ml) was collected at a concentration of 5.65 mg/ml and 300 μl were directly taken frozen for EPR spectroscopy of Fsr in its oxidized form.

Samples from the same methylene blue treated Fsr (passed through a Sephadex G-25M column) at 5.09 mg/ml final concentration were incubated for 5 minutes with 10 mM Na_2_SO_3_, and then stored in liquid nitrogen.

All EPR spectra were recorded on a Bruker Elexsys E580 X band spectrometer (digitally upgraded) with a 4122HQE cavity linked to an ESR 900 Oxford Instruments helium flow cryostat. Cryocooling was performed by a Stinger (Cold Edge Technologies) closed-cycle cryostat driven by an F-70 Sumitomo helium compressor. Our local glassblower produced EPR tubes from Ilmasil PN tubing (OD 4.7 mm and 0.5 mm wall thickness, Qsil, Langewiesen, Germany). Prior to use, the tubes were extensively cleaned with pipe cleaners to remove inadvertent contaminants. EPR spectra were simulated with Easyspin (51). The concentration of Fsr for the spin integration (using a 1 mM Cu^2+^-EDTA solution as standard) was obtained by dividing the Fe-concentration as determined with the ferene method (32) by 24, since siroheme does not release Fe. Fitting to the Nernst equation was performed in Excel.

### High resolution Clear Native PAGE (hrCN PAGE)

To visualize the expression levels of Fsr in HS^-^ versus SO_3_^2-^ grown cultures and to estimate the oligomerization of Fsr, hrCN PAGE was performed. 10 ml *of M. thermolithotrophicus* and *M. jannaschii* cultures, with either 2 mM HS^-^ or 2 mM SO_3_^2-^ as sulfur substrates, were grown for one night at 65 °C, standing. Cells were harvested by anaerobic centrifugation at 6,000 x *g* for 20 min at room temperature and the cell pellets were resuspended in 2 ml of 50 mM Tricine/NaOH pH 8.0 and 2 mM dithiothreitol. The cells were sonicated 4 x 70 % intensity for 10 seconds, followed by a 30 second break (MS 73 probe, SONOPULS Bandelin). The hrCN PAGE was run anaerobically and the protocol was adapted from Lemaire et al. (2018 (52)). The detailed procedure of hrCN running conditions can be found in the supporting information.

## Data Availability Statement

The structures were deposited in the protein data bank under the ID: 7NP8 for *Mj*Fsr and 7NPA for *Mt*Fsr.

## Acknowledgements

We thank the Max Planck Institute for Marine Microbiology and the Max Planck Society for continuous support. We are grateful for the genome sequencing performed in the Max Planck-Genome-center (Cologne) with specific regards to Dr. Bruno Huettel. We thank Jessica Soares for the EPR data acquisition and measurement of the Fe content. We acknowledge the SOLEIL synchrotron for beam time allocation and the beamline staff of Proxima-1 for assistance with data collection. Furthermore, we thank the staff of beamline PXIII from SLS and P11 at PETRA III. We are also thankful to Christina Probian and Ramona Appel for their continuous support in the Microbial Metabolism laboratory and Dr. Olivier Lemaire for his support. We thank Brenna L. Boehman for her deep reviewing of the manuscript.

## Author contributions

MJ cultivated both methanogens, purified and crystallized both Fsr. MJ performed all biochemical characterization. MJ and TW collected X-ray data and solved the structures. MJ refined both Fsr models and MJ with TW validated the models. MJ and AJP performed the redox titration experiments, and AJP the spectroscopic analyses. TW designed the research. All co-authors contributed to the writing of the article.

## Abbreviations

Fsr: F_420_-dependent sulfite reductase
Sir: sulfite reductase
dSir: dissimilatory sulfite reductase
aSir: assimilatory sulfite reductase
SDS/hrCN PAGE: Sodium dodecylsulfate / high-resolution clear native polyacrylamide gel electrophoresis
MCR: Methyl-coenzyme M reductase
Frh: F_420_-reducing [NiFe]-hydrogenase
EPR: Electron paramagnetic resonance
NHE: Normal hydrogen electrode

