## Supplemental information for "The structure of the F_420_-dependent sulfite-detoxifying enzyme from *Methanococcales* reveals a prototypical sulfite-reductase with assimilatory traits"

### Materials and methods.

### Sulfur-free cultivation medium for *Methanococcales*

Per liter of medium: 558 mg of $KH_2PO_4$ (final concentration 4.1 mM), 1 g of KCl (13.4 mM), 25.13 g of NaCl (430 mM), 840 mg of $NaHCO_3$ (10 mM), 37.5 mg of $CaCl_2 \cdot 2\ H_2O$ (2.5 mM), 7.725 g of $MgCl_2 \cdot 6\ H_2O$ (38 mM), 1.18 g of $NH_4Cl$ (22.06 mM), 61.16 mg of nitrilotriacetic acid (0.32 mM), 6.16 mg of $FeCl_2 \cdot 4\ H_2O$ (0.031 mM), 10 µl of 2 mM $Na_2SeO_3 \cdot 5\ H_2O$ stock (0.02 µM), 3.3 mg of $Na_2WO_4 \cdot 2\ H_2O$ (0.01 mM) and 2.42 mg of $Na_2MoO_4 \cdot 2\ H_2O$ (0.01 mM) were dissolved under constant stirring in a measuring cylinder with 750 ml of deionized $H_2O$ ($dH_2O$) (1). Resazurin (1 ml, 1.5 mM) was added (0.0015 mM) and 10 ml of sulfur-free trace elements (see below) were added subsequently. For *M. jannaschii*, 30.24 g PIPES (100 mM final) was used as a buffer and a pH 7.0 was adjusted using sodium hydroxide pellets. For *M. thermolithotrophicus* the pH was set to either 7.6 with 50 mM Tris/HCl as buffer or to 6.2 with 50 mM MES. The media was filled up to a final volume of 1 L by the addition of $dH_2O$.

The cultivation media was transferred in a 1 L pressure protected DURAN® laboratory bottle with a magnetic stirring bar. The Duran flask was closed with a butyl rubber stopper and degassed by applying 3 min of evacuation, followed by 30 seconds of ventilation with $1 \times 10^5$ Pa $N_2$ atmosphere, under constant magnetic stirring. This was repeated 15 times and at the final ventilation step an overpressure of $0.3 \times 10^5$ Pa $N_2$ was applied.

**Trace element composition:** A 100-fold-concentrated trace element solution was prepared by first dissolving 1.36 g of nitrilotriacetic acid (7.1 mM) in 800 ml $dH_2O$ under magnetic stirring. The pH was shifted to 6.2 by adding NaOH pellets. 89.32 mg of $MnCl_2 \cdot 6\ H_2O$ (0.45 mM), 183.3 mg of $FeCl_3 \cdot 6\ H_2O$ (0.68 mM), 60.27 mg of $CaCl_2 \cdot 2\ H_2O$ (0.41 mM), 180.8 mg of $CoCl_2 \cdot 6\ H_2O$ (0.76 mM), 90 mg of $ZnCl_2$ (0.66 mM), 37.64 mg of $CuCl_2$ (0.28 mM), 46 mg of $Na_2MoO_4 \cdot 2\ H_2O$ (0.19 mM), 90 mg of $NiCl_2 \cdot 6\ H_2O$ (0.38 mM) and 30 mg of $VCl_3$ (0.19 mM) was added separately. The trace element mixture was filled up to a final volume of 1 L with $dH_2O$.

**Growth of *M. jannaschii*.** Duran bottles (10 x 1 L) were sealed with butyl rubber stoppers and the gas phase was exchanged for $H_2/CO_2$ (80:20, $1 \times 10^5$ Pa). 100 ml of anaerobic cultivation media was transferred into each bottle (ratio of 1:10 of medium/gas phase), with 1 mM $Na_2SO_3$ as a sole sulfur source. 5 ml of overnight culture ($OD_{600nm}$: 0.9) was used as an inoculum for 100 ml media. No additional reductant was added. The cultures were placed at 65 °C, standing for at least one hour followed by an overnight shaking at 180 rotation per minute without light. The cells were harvested in exponential phase with a final $OD_{600nm}$ of 1.83, by immediately transferring them into an anaerobic tent ($N_2/CO_2$ atmosphere at a ratio of 90:10), followed by anaerobic centrifugation for 30 min at 6,000 x *g* at 4 °C. The cell pellet was transferred in a sealed bottle gassed with 0.3 X $10^5$ Pa $N_2$ and flash frozen in liquid $N_2$ to be stored at -80 °C.

**Growth of *M. thermolithotrophicus* for Fsr crystallization.** *M. thermolithotrophicus* was grown in a fermenter at 50 °C with 10 mM $SO_4^{2-}$ as a sole sulfur substrate. Since $SO_3^{2-}$ is an inevitable intermediate in the $SO_4^{2-}$ reduction pathway it requires the expression of Fsr. Therefore, 1.5 L of anaerobic cultivation medium with 10 mM $SO_4^{2-}$ were continuously bubbled with $H_2$ and $CO_2$ (80:20, 2 X $10^4$ Pa) and inoculated with 100 ml preculture ($OD_{600nm}$: 4.2). Since the fermenter is an open system, we set a more alkaline pH (7.6) to prevent evaporation of produced $S^{2-}$. Here, it should predominantly be present in form of $HS^-$ and not $H_2S$ and therefore stay for longer time in the medium. The pH was checked every two hours by using a pH indicator. The cells were grown until late exponential phase ($OD_{600nm}$ of 2.97) and then immediately transferred in an anaerobic tent ($N_2/CO_2$ atmosphere at a ratio of 90:10). Cells were harvested by anaerobic centrifugation for 30 min at 6,000 x *g* at 4 °C. 1.5 L of culture with an $OD_{600nm}$ of 2.97 yielded 19.25 g of cells (wet weight). The cell pellet was transferred in a sealed bottle, gassed with 0.3 X $10^5$ Pa $N_2$, flash frozen in liquid $N_2$ and stored at -80 °C.

**Growth of *M. thermolithotrophicus* for Fsr activity assays.** To perform enzymatic activity assays *M. thermolithotrophicus* was directly grown on 2 mM $SO_3^{2-}$. 10 x 1 L Duran bottles were sealed with butyl rubber stoppers and the gas phase was exchanged for $H_2$ and $CO_2$ (80:20, 1 X $10^5$ Pa). 100 ml of anaerobic cultivation media containing 50 mM MES at pH 6.2 was transferred into each bottle (ratio of 1:10 of medium/gas phase), with 2 mM $Na_2SO_3$ final as a sole sulfur source. 5 ml of overnight grown culture ($OD_{600nm}$ = 1.7) was used as an inoculum for 100 ml media. No additional reductant was added. The cultures were placed at 65 °C, standing overnight. The cells were grown until early exponential phase ($OD_{600nm}$ of 0.8) since we assumed that most $SO_3^{2-}$ has not been converted into $HS^-$ yet and that Fsr should be highly expressed and active. The cells were immediately harvested by transferring them in an anaerobic tent ($N_2/CO_2$ atmosphere at a ratio of 90:10), followed by anaerobic centrifugation for 30 min at 6,000 x *g* at 4 °C. The cell pellet was transferred in a sealed bottle, gassed with 0.3 X $10^5$ Pa $N_2$, flash frozen in liquid $N_2$ and stored at -80 °C.

**Sulfite growth inhibition.** *M. thermolithotrophicus* was grown on different $SO_3^{2-}$ concentrations to determine the growth-inhibiting threshold. For this, 250 ml serum flasks were sealed with a butyl rubber stopper and the gas phase was exchanged for $H_2$ and $CO_2$ (80:20, 1 X $10^5$ Pa). 10 ml of anaerobic cultivation media with a pH set at 6.2 with 50 mM MES was transferred into each bottle. Then different $Na_2SO_3$ concentrations (2 mM, 10 mM, 20 mM, 30 mM and 40 mM final) were added in triplicates as a sole sulfur source and 2 mM $Na_2S$ was used as a control. The cultures grew at 65 °C for 22 hours, standing.

**Growth of *M. thermolithotrophicus* for titrations and EPR spectroscopy.** Due to the high demand of *Mt*Fsr for titration and EPR spectroscopy experiments *M. thermolithotrophicus* was grown in a 10 L fermenter with $SO_4^{2-}$ as a sole sulfur substrate and another 10 L fermenter with $SO_3^{2-}$ as a sole sulfur source to boost *Mt*Fsr natural expression. The fermenter containing $SO_4^{2-}$ was performed as mentioned above with an inoculum of 350 ml ($OD_{600nm}$: 3.2). 7.4 L of culture with an $OD_{600nm}$ of 4.8 yielded 74 g of cells (wet weight). In the $SO_3^{2-}$ fermenter, *M. thermolithotrophicus* was grown at 50 °C in 7 L anaerobic cultivation medium with a pH of 6.2 supplemented with 5 mM $SO_3^{2-}$ as a sole sulfur substrate, continuously bubbled with $H_2$ and $CO_2$ (80:20, 2 X $10^4$ Pa). 600 ml preculture ($OD_{600nm}$: 2.34) were used as inoculum. The cells were grown until an $OD_{600nm}$ of 2.48 and then immediately transferred in an anaerobic tent ($N_2/CO_2$ atmosphere at a ratio of 90:10). Cells were harvested by anaerobic centrifugation for 30 min at 6,000 x *g* at 4 °C and a final yield of 51 g of cells (wet weight) was obtained.

The cell pellets were transferred in a sealed bottle, gassed with 0.3 X $10^5$ Pa $N_2$, flash frozen in liquid $N_2$ and stored at -80 °C.

**Genome sequencing of *M. thermolithotrophicus*.** *M. thermolithotrophicus* was anaerobically grown in the above described medium and 2 mM sodium sulfide was used as a sulfur source. A total culture volume of 20 ml was used. Cells were aerobically harvested by centrifugation (30 min, 6,000 x *g* at 4 °C). DNA was extracted and purified based on Martín-Platero et al. 2007 (2). Quality control, library preparation and sequencing (PacBio Sequel II) were performed in the Max Planck-Genome center (Cologne).

**Purification of *Mj*Fsr.** *M. jannaschii* cells (13.5 g wet weight) were thawed under warm water and transferred in an anaerobic tent ($N_2/CO_2$ atmosphere at a ratio of 90:10). Cells were diluted by three volumes of lysis buffer (50 mM Tricine/NaOH pH 8.0, 2 mM dithiothreitol (DTT)) and disrupted by sonication: 7 cycles at 62 % intensity with 30 pulses followed by 1 min break (probe MS76, SONOPULS Bandelin). Cell debris were removed anaerobically via centrifugation (21,000 x *g*, one hour, room temperature). The protein concentration (measured by Bradford) of the supernatant was estimated to 4.68 $mg.ml^{-1}$. The supernatant was transferred to a Coy tent ($N_2/H_2$ atmosphere of 97:3) under yellow light at 20 °C. The sample was diluted with two volumes of lysis buffer and passed through a 0.2 µm filter (Sartorius). The filtered sample was loaded on a 10 ml Q-sepharose high-performance column (GE Healthcare) which was previously equilibrated with 5 column volumes (CV) of lysis buffer. The column was then washed with 2 CV of lysis buffer. Fsr was eluted by a gradient of NaCl (from 0.1 to 0.6 M) in 27 CV at a flow rate of 1.5 $ml.min^{-1}$ in fraction sizes of 3.5 ml. *Mj*Fsr eluted between 0.37 to 0.41 M NaCl. The fractions of interest were pooled and 1:1 diluted with HIC buffer (25 mM Tris/HCl pH 7.6, 2 M $(NH_4)_2SO_4$ and 2 mM DTT). The sample was filtered and applied to a Source15Phe 4.6/100 PE column (GE healthcare) previously equilibrated with the HIC buffer. The column was then washed with 2 CV of 25 mM Tris/HCl pH 7.6, 1.4 M $(NH_4)_2SO_4$ and 2 mM DTT buffer. The elution was performed at a flow rate of 0.8 $ml.min^{-1}$ by a decreasing gradient of $(NH_4)_2SO_4$ (1.4 to 0 M) over 90 min, with a fractionation size of 2 ml. Fsr eluted in the fractions at 0.9 to 0.78 M $(NH_4)_2SO_4$. Those fractions were merged and concentrated using a 30-kDa-cutoff filter (Merck Millipore, Darmstadt, Germany). The concentrated sample was passed through 0.2-µm filter and injected on a Superdex 200 Increase 10/300 GL (GE Healthcare) equilibrated in storage buffer (25 mM Tris/HCl pH 7.6, containing 10 % v/v glycerol and 2 mM DTT). The elution was performed at a flow rate of 0.4 $ml.min^{-1}$ in the storage buffer. *Mj*Fsr eluted as a sharp Gaussian peak at 10.4 ml. The pooled samples were concentrated by passing them through a 30-kDa-cutoff filter, and the final concentration was measured by the Bradford method (Bio-Rad, Munich, Germany). The sample was immediately crystallized at a concentration of 6.1 $mg.ml^{-1}$.

**Purification of *Mt*Fsr for crystallization.** Cells (19.25 g wet weight) derived from a fermenter were thawed under warm water and transferred to an anaerobic tent containing an atmosphere of $N_2/CO_2$ (90:10). Cells were lysed by osmotic shock through the addition of 60 ml lysis buffer (50 mM Tricine/NaOH pH 8.0, 2 mM DTT). Cell lysate was homogenized by sonication: 3 cycles at 70 % intensity with 30 pulses followed by 1 min break (probe MS76, SONOPULS Bandelin) and cell debris were removed anaerobically via centrifugation (21,000 x *g*, one hour at 4 °C). The supernatant was transferred in a Coy tent ($N_2/H_2$ atmosphere of 97:3), with yellow light at 20 °C. The sample was filtered through a 0.2 µm filter (Sartorius) and was passed onto a DEAE fast flow column (30 ml), equilibrated with lysis buffer. The column was then washed with 2 CV of lysis buffer. *Mt*Fsr was eluted with a gradient of 0.1 to 0.6 M NaCl in 120 min at a flow rate of 2.5 $ml.min^{-1}$ and in fractionation sizes of 4 ml. *Mt*Fsr eluted between 0.3 to 0.39 M NaCl. The fractions of interest were merged, diluted by 3 volume of lysis buffer and filtered through a 0.2 µm filter. The filtered sample was loaded on a 15 ml Q Sepharose high performance column, equilibrated with lysis buffer. The column was washed with 2 CV of lysis buffer. A gradient of 0.15 to 0.55 M NaCl in 120 min with a flow rate of 1 $ml.min^{-1}$ was performed and fractions of 1.5 ml were collected. *Mt*Fsr eluted between 0.49 to 0.53 M NaCl. Fractions of interest were pooled and diluted with 2 volumes of HAP buffer (20 mM $K_2HPO_4$/HCl pH 7.0 and 2 mM DTT) and subsequently filtered through a 0.2 µm filter. The filtered sample was applied to a 10 ml hydroxyapatite column type 1 (Bio-Scale Mini CHT cartridges, BioRad) equilibrated with HAP buffer. The column was washed with 2 CV of HAP buffer and a gradient of 0.02 to 0.5 M $K_2HPO_4$ for 60 min at a flow rate of 2 $ml.min^{-1}$ was performed and 3 ml fractions were collected. *Mt*Fsr eluted between 0.28 to 0.39 M $K_2HPO_4$ and the respective fractions were pooled. The pool was diluted 1:3 with 25 mM Tris/HCl pH 7.6, 2 M $(NH_4)_2SO_4$ and 2 mM DTT (HIC Buffer). The filtered sample was applied to a Source15Phe 4.6/100 PE column (GE healthcare) previously equilibrated with the HIC buffer. The column was then washed with 2 CV of HIC buffer. A gradient $(NH_4)_2SO_4$ ranging from 2 to 1 M was performed for 30 min at a flow rate of 0.8 $ml.min^{-1}$ with a fractionation size of 1 ml. *Mt*Fsr eluted between 1.38 to 1.23 M $(NH_4)_2SO_4$ and the respective fractions were pooled. The buffer was exchanged for the storage buffer (25 mM Tris/HCl pH 7.6, containing 10 % v/v glycerol and 2 mM DTT) by using 30-kDa-cutoff filter (6 ml, Merck Millipore, Darmstadt, Germany) and *Mt*Fsr was concentrated to 11.06 mg. $ml^{-1}$ in a volume of 120 µl. The protein concentration was estimated by the Bradford method. The sample was immediately crystallized.

**Purification of *Mt*Fsr for enzyme activity assays.** $SO_3^{2-}$-grown cells (8 g wet weight) were thawed under warm water and transferred to an anaerobic tent containing an atmosphere of $N_2/CO_2$ (90:10). Cells were lysed by osmotic shock through the addition of 60 ml lysis buffer (50 mM Tricine/NaOH pH 8.0, 2 mM DTT). Cell lysate was homogenized by sonication: 9 cycles at 75 % intensity with 30 pulses followed by 1 min break (probe KE76, SONOPULS Bandelin) and cell debris were removed anaerobically via centrifugation (21,000 x *g*, one hour at 4 °C). The supernatant was transferred to a Coy tent ($N_2/H_2$ atmosphere of 97:3) under yellow light at 20 °C and was diluted with 90 ml lysis buffer, filtered through a 0.2 µm filter. The filtered sample was applied to a 10 ml DEAE fast flow column (GE healthcare), which was previously equilibrated with lysis buffer. The column was then washed with 2 CV of lysis buffer. A gradient of 0.1 to 0.6 M NaCl was applied for 120 min at a flow rate of 2.5 $ml.min^{-1}$ and fractions of 4 ml were collected. *Mt*Fsr eluted between 0.34 to 0.4 M NaCl. The fractions of interest were merged and diluted by 3 volumes of lysis buffer. The filtered sample was loaded on a 10 ml Q Sepharose high performance column (GE healthcare) and a gradient of 0.15 to 0.55 M NaCl was applied for 120 min with a flow rate of 1 $ml.min^{-1}$. Fractions of 1.5 ml were collected. *Mt*Fsr eluted between 0.49 to 0.53 M NaCl. The *Mt*Fsr fractions were pooled, and three times diluted with HAP buffer (20 mM $K_2HPO_4$/HCl pH 7.0 and 2 mM DTT). The filtered sample was applied to a 10 ml hydroxyapatite type 1 (Bio-Scale Mini CHT cartridges, BioRad) equilibrated with HAP buffer. The column was washed with 2 CV of HAP buffer and a gradient of 0.02 to 0.5 M $K_2HPO_4$ in 60 min at a flow rate of 2 $ml.min^{-1}$ was performed. 1.5 ml fractions were collected. *Mt*Fsr eluted between 0.25 to 0.42 M $K_2HPO_4$ and the respective fractions were pooled. The pool was diluted with 3 volumes of HIC buffer (25 mM Tris/HCl pH 7.6, 2 M $(NH_4)_2SO_4$ and 2 mM DTT). The filtered sample was applied to a Source15Phe 4.6/100 PE column (GE healthcare) previously equilibrated with the HIC buffer. The column was then washed with 2 CV of 25 mM Tris/HCl pH 7.6, 1.6 M $(NH_4)_2SO_4$ and 2 mM DTT buffer. *Mt*Fsr was eluted in a gradient of 1.6 to 0.8 M of $(NH_4)_2SO_4$ in 25 min at a flow rate of 0.8 $ml.min^{-1}$ and a fractionation size of 1 ml. *Mt*Fsr eluted between 1.43 to 1.28 M $(NH_4)_2SO_4$ and the respective fractions were pooled. The buffer was exchanged for the storage buffer (25 mM Tris/HCl pH 7.6, containing 10 % v/v glycerol and 2 mM DTT) by using 30-kDa-cutoff filter (6 ml, Merck Millipore, Darmstadt, Germany) and *Mt*Fsr was concentrated to 900 µl. The concentrated sample was passed onto a Superdex 200 Increase 10/300 GL (GE Healthcare), equilibrated in storage buffer. *Mt*Fsr eluted at a flow rate 0.4 $ml.min^{-1}$ in a sharp Gaussian peak at an elution volume of 10.01 ml (Fig. S2). To determine the apparent molecular weight of *Mt*Fsr, standard proteins (Conalbumin, Aldolase and Ferritin, purchased from GE Healthcare) were passed at the same flow rate and in the same buffer. The fractions of interest containing *Mt*Fsr were concentrated with a 30-kDa cut-off centrifugal concentrator to 1 ml and the protein was directly used for enzymatic activity assays. The concentration of purified *Mt*Fsr, estimated by the Bradford method, was 3.41 $mg.ml^{-1}$.

**Purification of *Mt*Fsr for titrations and EPR spectroscopy.** For the titrations and EPR spectroscopic measurements two separate purifications were carried out starting either with 34 g cells (wet weight) derived from a $SO_3^{2-}$-grown fermenter or with 49.5 g cells (wet weight) derived from a $SO_4^{2-}$-grown fermenter. Cells were thawed under warm water and transferred to an anaerobic tent containing an atmosphere of $N_2/CO_2$ (90:10). Cells were lysed by osmotic shock through the addition of 180 ml and 240 ml lysis buffer (50 mM Tricine/NaOH pH 8.0, 2 mM DTT), respectively. The cell lysates were homogenized by sonication: 4 cycles at 72 % intensity with 60 pulses followed by 1.30 minutes break (probe MS76, SONOPULS Bandelin) and the cell debris were removed anaerobically via centrifugation (21,000 x g, 1 h at 10 °C). The supernatant was transferred into a Coy tent ($N_2/H_2$ atmosphere of 97:3), with yellow light at 20 °C.
The following purification steps were carried out as described in "Purification of *Mt*Fsr for crystallization". In the final purification step the buffer was exchanged by dilution and concentration into storage buffer (25 mM Tris/HCl pH 7.6, containing 10 % v/v glycerol and 2 mM DTT) by using 30-kDa-cutoff filter (6 ml, Merck Millipore, Darmstadt, Germany). *Mt*Fsr derived from the $SO_3^{2-}$-grown fermenter was concentrated to 18 $mg.ml^{-1}$ in a volume of 4.54 ml and for the $SO_4^{2-}$-grown fermenter MtFsr was concentrated to 20 $mg.ml^{-1}$ in a volume of 1.24 ml. The protein concentrations were estimated by the Bradford method.

**Purification of the $F_{420}$-reducing hydrogenase (Frh) from *Methanothermococcus thermolithotrophicus*.** Frh was required to reduce $F_{420}$ and was purified from the same batch of cells as *Mt*Fsr, used for crystallization. The activity of *Mt*Frh after each purification step was followed by the reduction of methyl-viologen in the $N_2/H_2$ tent (97:3). The assay was performed in 120 µl of 0.5 M $KH_2PO_4$/NaOH pH 7.6 containing 1.7 mM of oxidized methyl-viologen. The addition of 2 µl from the fractions containing Frh led to a blue coloration.

*Mt*Frh was in the same pool as *Mt*Fsr, used for crystallization, for the DEAE- and the Q-sepharose column. The Q-Sepharose column performed the separation of the two target proteins. *Mt*Frh eluted between 0.48 to 0.49 M NaCl from the Q-Sepharose column. The filtered sample was applied to a 10 ml hydroxyapatite type 1 (Bio-Scale Mini CHT cartridges, BioRad) equilibrated with HAP buffer (20 mM $K_2HPO_4$/HCl pH 7.0 and 2 mM DTT). The column was then washed with 2 CV of HAP buffer. The elution was performed with a gradient of 0.02 to 0.5 M $K_2HPO_4$ in 60 min at a flow rate of 2 $ml.min^{-1}$ with 3 ml fractions. *Mt*Frh eluted between 0.22 to 0.37 M $K_2HPO_4$ and the respective fractions were pooled. The pool was diluted 1:1 with the HIC buffer (25 mM Tris/HCl pH 7.6, 2 M $(NH_4)_2SO_4$ and 2 mM DTT). The filtered sample was applied onto a Source15Phe 4.6/100 PE column (GE Healthcare) previously equilibrated with the HIC buffer. The column was then washed with 2 CV of 25 mM Tris/HCl pH 7.6, 1.0 M $(NH_4)_2SO_4$ and 2 mM DTT buffer. *Mt*Frh was eluted in a gradient of 1 to 0 M $(NH_4)_2SO_4$ in 30 min at a flow rate of 0.8 $ml.min^{-1}$ and a fractionation size of 1 ml. *Mt*Frh eluted between 0.4 to 0.15 M $(NH_4)_2SO_4$ and the respective fractions were pooled. The buffer was exchanged for the storage buffer (25 mM Tris/HCl pH 7.6, containing 10 % v/v glycerol and 2 mM DTT) by using 30-kDa-cutoff filter (6 ml, Merck Millipore, Darmstadt, Germany) and *Mt*Frh was concentrated to 4.97 mg. $ml^{-1}$ in 100 µl. The purified sample was aliquoted and anaerobically flash frozen in liquid $N_2$ and stored at -80 °C. *Mt*Frh lost its activity after more than 1 cycle of thawing-freezing.

**Purification of oxidized $F_{420}$.** Since $F_{420}$ is highly sensitive to light, all steps were carried out under yellow light or by covering the sample with aluminum foil. About ~10 g (wet weight) of *M. thermolithotrophicus* cells from a 1.5 L fermenter were anaerobically lysed by osmotic shock and sonication (see above). The sample was centrifuged at 45,000 x *g* for 60 min at 4°C. The supernatant was transferred in a Coy tent containing an atmosphere of $N_2/H_2$ (97:3). The sample was filtered and passed onto a 30 ml DEAE sepharose column equilibrated with 50 mM Tricine/NaOH pH 8.0 and 2 mM DTT. $F_{420}$ was eluted by a gradient of 0 to 0.6 M NaCl. The samples containing $F_{420}$ were determined based on the absorption profile at 420 nm and eluted between 0.48 M and 0.58 M NaCl. Pooled fractions were moved outside of the tent and diluted with one volume of HIC-$F_{420}$ buffer (25 mM Tris/HCl pH 7.6, 2 M $(NH_4)_2SO_4$)). $(NH_4)_2SO_4$ powder was directly added to the diluted sample to reach a final concentration of 3 M $(NH_4)_2SO_4$ and was stirred for one hour at room temperature. The sample was centrifuged at 4,000 x *g* for 20 minutes at room temperature. The supernatant was filtered through a 0.2 µm filter and loaded on a 30 ml Phenyl-Sepharose high-performance column (30 ml), equilibrated with HIC-$F_{420}$ buffer. $F_{420}$ was eluted with a gradient of 2 to 0 M $(NH_4)_2SO_4$ in 20 min, at a flow rate of 2 $ml.min^{-1}$ and 1 ml fractions. The fractions containing $F_{420}$ were pooled and filtered through a 0.2 µm filter. The sample was diluted by 50 volumes of 5 mM Tris/HCl pH 8.0 and loaded overnight on a 5 ml Q-Sepharose high-performance column, equilibrated in 5 mM Tris-HCl pH 8.0. The following steps were performed at 4 °C. The column containing the bound $F_{420}$ was washed with 5 CV of 20 mM $(NH_4)HCO_3$ pre-cooled at 4°C. $F_{420}$ elution was performed by adding 1 M $(NH_4)HCO_3$ and collected in a brown serum flask. $(NH_4)HCO_3$ was removed by evacuation at 37°C for 2 hours under constant stirring. $(NH_4)HCO_3$ free $F_{420}$ powder was obtained by freeze drying. The purity of the preparation was checked by measuring the ratio of $Abs_{247}/Abs_{420}$ in 25 mM Tris pH 8.8. A pure sample would have a ratio value of 0.85 (3). $F_{420}$ concentration was estimated by measuring the absorbance at 420 nm in 25 mM Tris pH 7.5 ($\varepsilon_{420nm}$=41.4 $mM^{-1}.cm^{-1}$). The final concentration of oxidized $F_{420}$ used for this study was 3.15 mM.

**Reduction of $F_{420}$ for enzyme assays.** For enzyme activity assays and co-crystallization of *Mt*Fsr with $F_{420}H_2$, the oxidized $F_{420}$ needed to be reduced. Dithionite was not used since it might generate $SO_3^{2-}$ as a side reaction. All steps were performed under the strict exclusion of oxygen and under yellow light. First, the aerobic gas phase of the $F_{420}$-stock was exchanged several times for $N_2$. The sample was then transferred in a Coy tent with an atmosphere containing a $N_2/H_2$ mixture (97:3). The reduction took place in 1.4 ml of 200 mM $KH_2PO_4$, pH 7.0, 0.5 mM $F_{420}$ and 5 µl of 5 mg. $ml^{-1}$ purified *Mt*Frh was added. Outside of the tent, in a brown serum flask, the gas phase was exchanged three times for $H_2$ and $CO_2$, by evacuation and gassing with 1 X $10^5$ Pa $H_2$ and $CO_2$ (80:20) at room temperature. The reduction of $F_{420}$ was observed by the color shift from yellow to transparent. Frh was removed by passing the sample through a 10 kDa cut-off filter. Since reduced $F_{420}$ is not stable and oxidized with time, aliquoted $F_{420}H_2$ without Frh was immediately flash frozen in liquid $N_2$ and stored at – 80 °C.

**Reduction of $F_{420}$ for redox titrations.** $F_{420}$ is the physiological electron donor for Fsr and was therefore used as the reductant for the redox titrations. Oxidized $F_{420}$ was purified as previously described. Since both, the reduction of $F_{420}$ with Frh is not complete, and $F_{420}H_2$ is not stable over time, we reduced $F_{420}$ with sodium borohydride, like previously described (4). The reduction of $F_{420}$ was performed in an anaerobic chamber with an $N_2/H_2$ atmosphere of 97:3 at 25 °C. $F_{420}H_2$ was generated by reducing 100 µl $F_{420}$ at 7.53 mM with a few sodium borohydride crystals in a 10 mM Tris/HCl solution at pH 7.6, followed by destruction of excess borohydride by acidification with 50 µl 1 M hydrochloric acid. After the hydrogen evolution ceased, the pH was readjusted by the addition of 50 µl 1 M Tris/HCl pH 8.0. The generated $F_{420}H_2$ was prepared freshly for each experiment and immediately used.

**High resolution Clear Native PAGE (hrCN PAGE).** Linear polyacrylamide gradient gels (8 to 15 %) were prepared under aerobic conditions but then transferred into an anoxic chamber (atmosphere of $N_2/CO_2$, 90:10), where the gels were equilibrated in anaerobic cathode buffer (50 mM Tricine; 15 mM Bis-Tris, pH 7; 0.05 % w/v sodium deoxycholate; 0.01 % w/v dodecyl maltoside and 2 mM of DTT) overnight. Fresh and anaerobic samples were diluted with the lysis buffer to a final concentration of 1 $mg.ml^{-1}$ and a volume of 12 µl per sample was loaded on the gel, as well as 2 µl of the NativeMark™ Unstained Protein Standard ladder (ThermoFisher). Glycerol (20 % v/v final) was added to each sample and 0.001 % w/v Ponceau S served as a marker for protein migration. The electrophoresis anode buffer contained 50 mM Bis-Tris buffer pH 7 and 2 mM DTT. The hrCN gels were run with a constant 40 mA current (PowerPac™ Basic Power Supply, Bio-Rad). After electrophoresis, the protein bands were aerobically stained with Instant Blue™ (Expedeon).

**Supplementary Table 1.** X-ray analysis statistics for Fsr.

| | ***Mj*Fsr SAD at the Fe-K edge** | ***Mj*Fsr** | ***Mt*Fsr** |
|---|---|---|---|
| **Data collection** | | | |
| Synchrotron source | SOLEIL, PX1 | SOLEIL, PX1 | SLS, PXIII |
| Wavelength (Å) | 1.74013 | 0.97857 | 1.00004 |
| Space group | $C222_1$ | $C222_1$ | $P1$ |
| Resolution (Å) | 120.05 – 2.32 (2.54 – 2.32) | 78.82 – 2.30 (2.41 – 2.30) | 121.28 – 1.55 (1.69 – 1.55) |
| Cell dimensions | | | |
| a, b, c (Å) | 167.34 172.34 196.01 | 167.26 172.20 195.89 | 113.15, 124.16, 241.06 |
| α, β, γ (°) | 90, 90, 90 | 90, 90, 90 | 102.28, 95.71, 90.25 |
| $R_{merge}$(%)[a] | 37.4 (260.1) | 25.1 (162.3) | 18.3 (162.3) |
| $R_{pim}$ (%)[a] | 10.3 (77.4) | 7.0 (45.9) | 7.5 (66.4) |
| $CC_{1/2}$ [a] | 0.996 (0.596) | 0.995 (0.629) | 0.996 (0.439) |
| $I/\sigma_I$[a] | 10.4 (1.5) | 8.3 (1.6) | 8.7 (1.5) |
| Spherical completeness[a] | 74.8 (15.9) | 83.2 (32.3) | 75.5 (16.6) |
| Ellipsoidal completeness[a] | 95.0 (66.7) | 96.0 (94.8) | 94.5 (70.9) |
| Redundancy[a] | 26.9 (22.7) | 13.9 (13.2) | 7.0 (6.9) |
| Nr. unique reflections[a] | 91281 (4565) | 104064 (5203) | 1396397 (69018) |
| **Refinement** | | | |
| Resolution (Å) | | 64.67 – 2.30 | 77.29 – 1.55 |
| Number of reflections | | 104036 | 1396186 |
| $R_{work}/R_{free}$[b] (%) | | 18.15/20.43 | 15.88/17.11 |
| Number of atoms | | | |
| Protein | | 19554 | 79372 |
| Ligands/ions | | 920 | 3550 |
| Solvent | | 772 | 10778 |
| Mean B-value (Å$^2$) | | 41.25 | 26.37 |
| Molprobity clash score, all atoms | | 1.64 | 3.22 |
| Ramachandran plot | | | |
| Favored regions (%) | | 98.42 | 97.65 |
| Outlier regions (%) | | 0 | 0 |
| rmsd[c] bond lengths (Å) | | 0.010 | 0.011 |
| rmsd[c] bond angles (°) | | 1.384 | 1.371 |
| **PDB ID code** | | 7NP8 | 7NPA |

[a] Values relative to the highest resolution shell are within parentheses. [b] $R_{free}$ was calculated as the $R_{work}$ for 5 % of the reflections that were not included in the refinement. [c] rmsd, root mean square deviation.

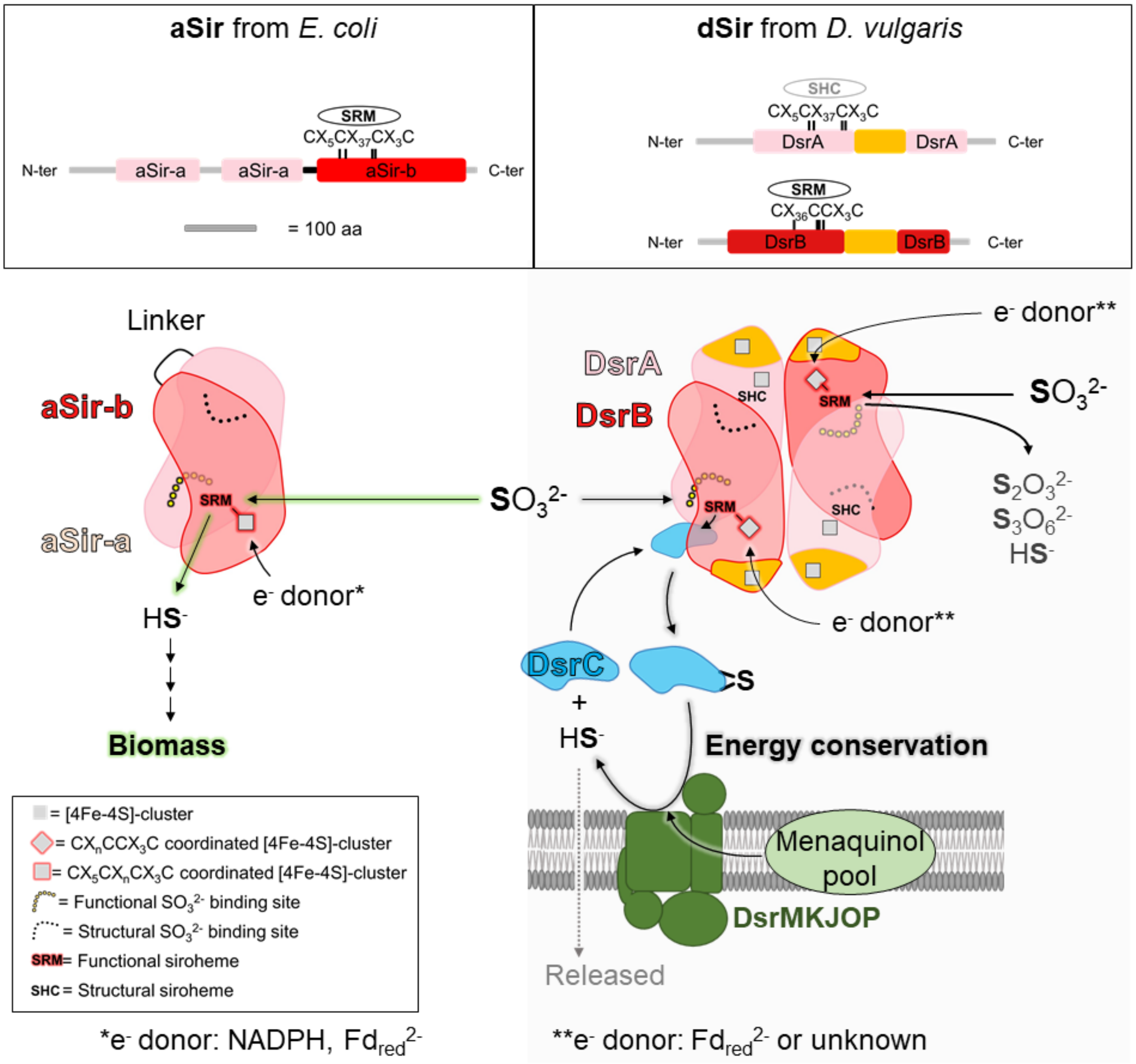


**Supplementary Figure S1.** Structural and functional organization of assimilatory (aSir) and dissimilatory (dSir refer here as DsrAB) sulfite reductases,. Top panel: visualization of distinct and conserved domains in aSir as well as dSir. The [4Fe-4S]-cluster binding motifs in the proximity of the siroheme or sirohydrochlorin are highlighted. The same scale was applied for all polypeptides. Bottom panel: aSirs (left) are functional monomers that probably evolved through a gene duplication event, where one gene lost its cluster binding motif. The N-terminal half abbreviated as aSir-a (colored in light pink) has a structural function and the C-terminal half abbreviated as aSir-b (red) harbors the active [4Fe-4S]-siroheme. aSirs indirectly use electrons from NADPH (bacteria) or directly via a [2Fe-2S]-cluster containing ferredoxin (plants) to reduce $SO_3^{2-}$ to $HS^-$ in a six-electron reduction reaction (5, 6). The produced sulfide will be used for Sulfur assimilation. dSirs (right) are composed of two DsrA (light pink) and two DsrB (red) subunits and receive electrons from reduced ferredoxins ($Fd_{red}^{2-}$) or so far unknown donors (7). In absence of DsrC (cyan), DsrAB reduces $SO_3^{2-}$ to thionates (like $S_2O_3^{2-}$, $S_3O_6^{2-}$) and $HS^-$. In presence of DsrC, the intermediate Sulfur species bound on the siroheme of DsrAB is transferred to DsrC. In the case of *Desulfovibrio* species, the membrane DsrMKJOP complex (green) fully reduces the DsrC-trisulfide (4 electrons transfer) probably by using the menaquinol pool and generates DsrC and $HS^-$ via the trisulfide pathway, a key process for energy conservation (7). The product $HS^-$ will be released into the environment and DsrC is recycled for the next reduction reaction.

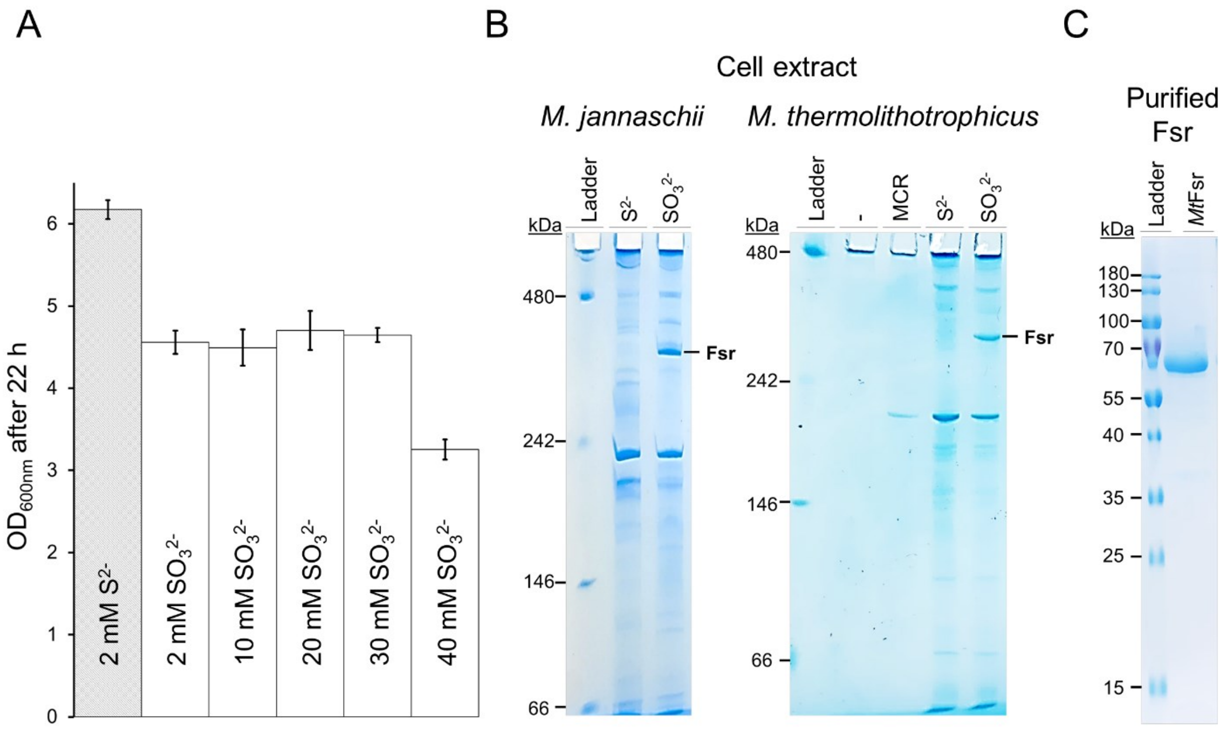


**Supplementary Figure S2.** Physiological and expression profiles of Fsr from *Methanococcales*. (A) Final $OD_{600nm}$ of *M. thermolithotrophicus* grown on sulfide ($S^{2-}$) and different sulfite ($SO_3^{2-}$) concentrations as a sole sulfur source after 22 hours (*N*=3). (B) hrCN-PAGE of cell extracts (12 µg loaded) from *M. jannaschii* (left) and *M. thermolithotrophicus* (right), grown on 2 mM $S^{2-}$ or 2 mM $SO_3^{2-}$ as a sole sulfur source. Purified MCR from *M. thermolithotrophicus* (1.7 µg loaded) was used as a control for the hrCN-PAGE (8). (C) SDS-PAGE profile of purified *Mt*Fsr.

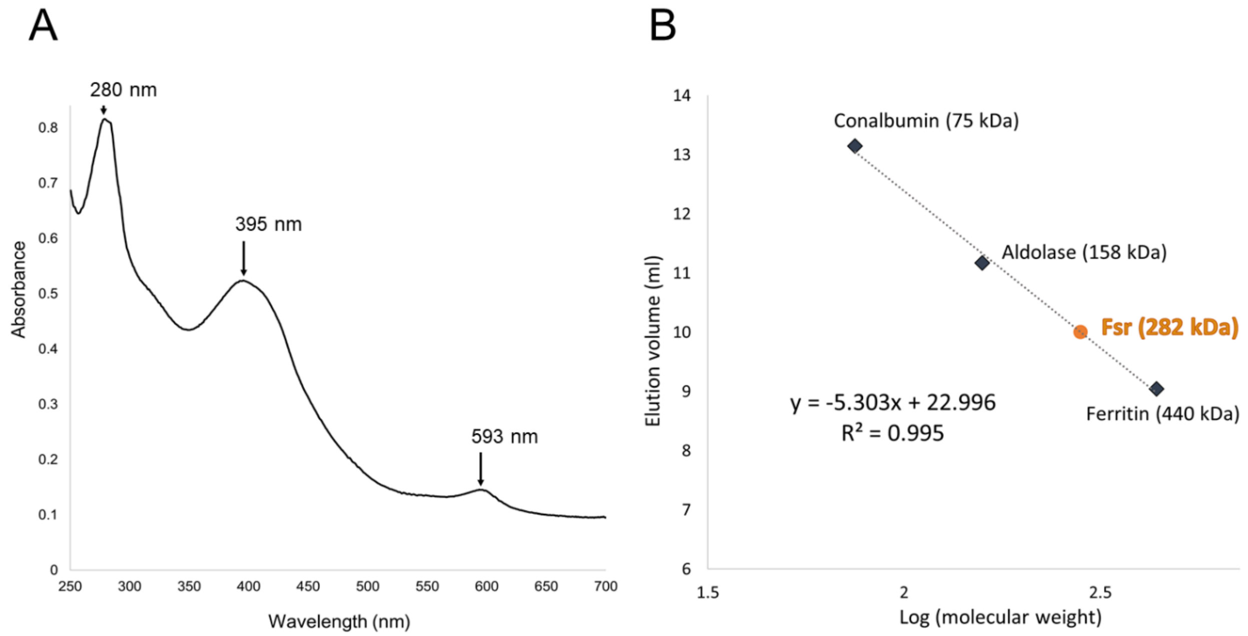


**Supplementary Figure S3.** (A) UV-visible spectrum of 0.33 mg of *Mt*Fsr anaerobically prepared (100 % $N_2$) measured in a total volume of 0.8 ml in 25 mM Tris/HCl, pH 7.6, 0.15 M NaCl, 10 % v/v glycerol and 2 mM DTT. *Mt*Fsr displays the same UV-visible spectra typical for [Fe-S]-cluster and siroheme containing enzymes (9). The UV spectrum of *Mj*Fsr was determined by Johnson and Mukhopadhyay and exhibited three peaks at 280 nm, 395 nm and 593 nm (10). (B) Molecular weight estimation of *Mt*Fsr via size exclusion chromatography (Superdex 200 Increase 10/300 GL from GE Healthcare). Apparent molecular weight of purified *Mt*Fsr was calculated based on the elution profile and estimated to 281.59 kDa. The theoretical weight of a monomeric *Mt*Fsr based on its amino acid sequence is 69.145 kDa. *Mt*Fsr is therefore apparently organized as a homotetramer (theoretical molecular weight of the protein in the homotetramer: 276.58 kDa).

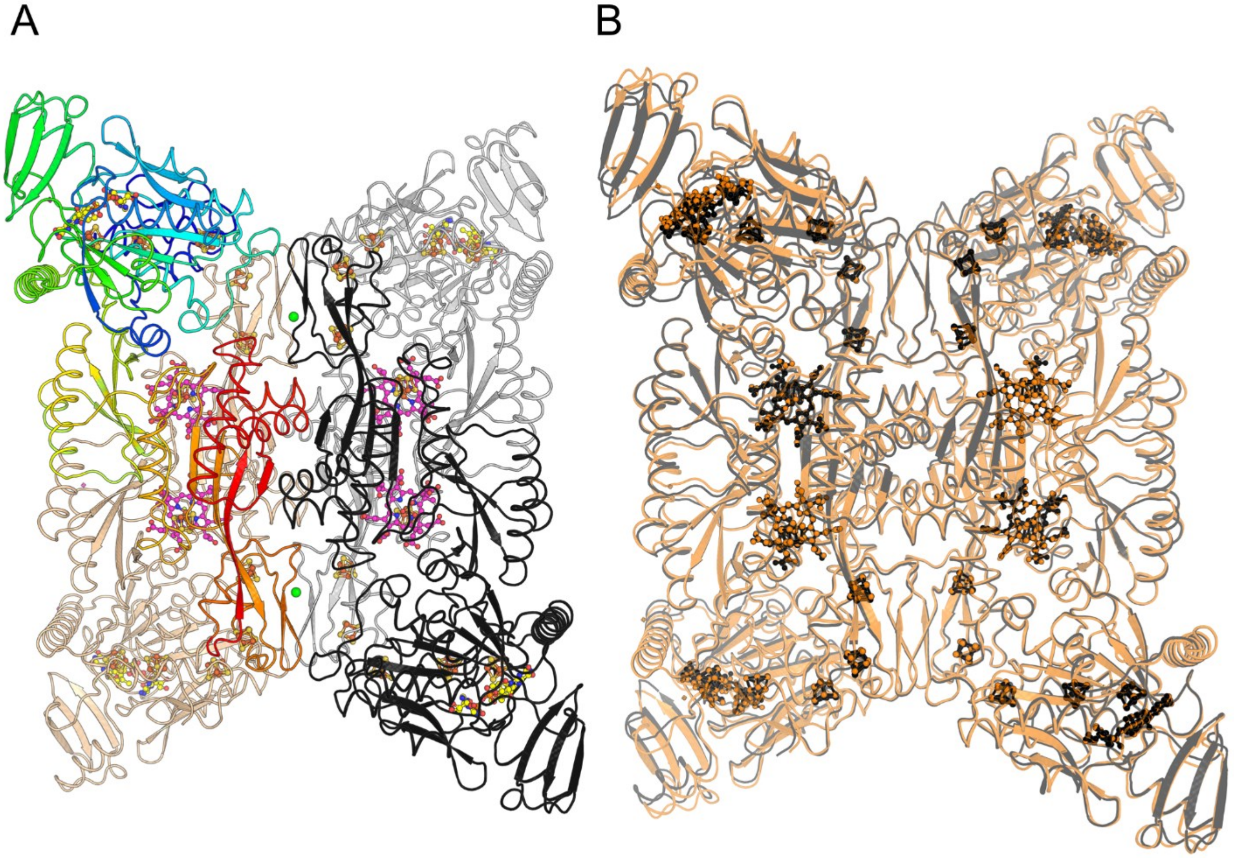


**Supplementary Figure S4.** Homotetrameric arrangement of Fsr. (A) General organization of *Mj*Fsr. The four monomers are shown in cartoon and colored in tan, black and white, with the last monomer colored in rainbow ranging from blue (N-terminal start, ferredoxin domain) to red (C-terminal end in the sulfite reductase domain). Ligands are shown as balls and sticks with nitrogen, oxygen, sulfur, and iron colored in blue, red, yellow, and orange. Carbons are colored in yellow for FAD and pink for siroheme. The green sphere placed at the inter-dimer interface corresponds to calcium ions coordinated by conserved aspartates and waters. (B) Superposition of *Mt*Fsr (black) with *Mj*Fsr (orange, rmsd of 0.456 Å for 544-Cα aligned). Ligands are shown in balls and sticks and colored in black and red for *Mt*Fsr and *Mj*Fsr, respectively.

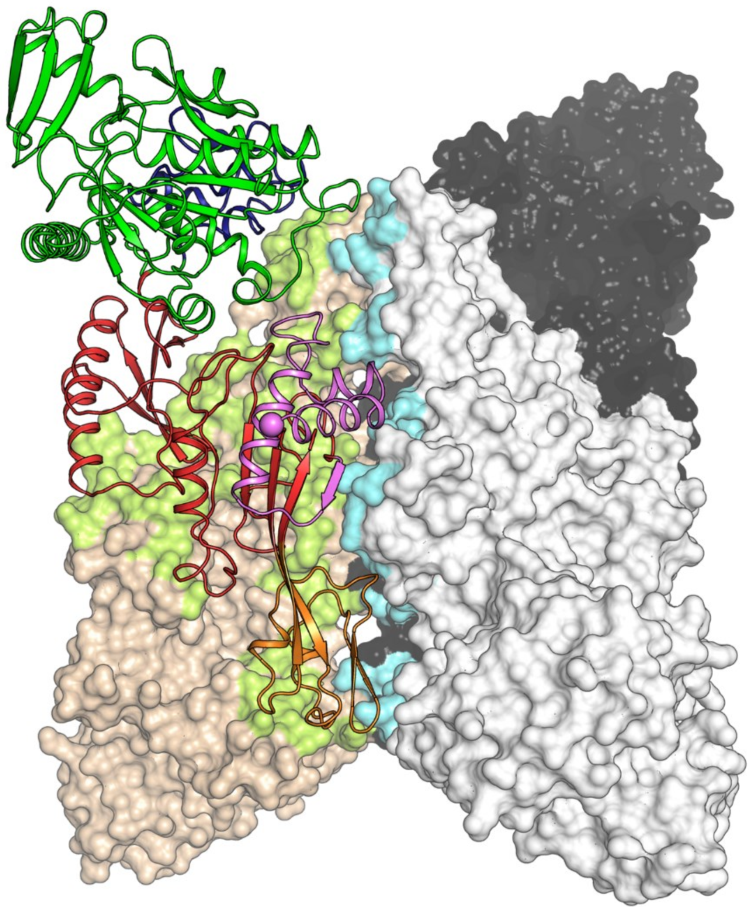

**Supplementary Figure S5.** Surface area involved in the oligomerization in Fsr. Monomers of *Mt*Fsr are shown in surface representation, with one monomer being displayed in cartoon and colored by its domain composition: N-terminus ferredoxin domain in dark blue, $F_{420}H_2$-oxidase in green, the sulfite reductase domain in red and its inserted ferredoxin domain in orange. The C-terminal segment involved in the oligomerization is colored in light pink with the C-terminus highlighted as a ball. The monomer-monomer contacts are colored in green surface and contacts to the adjacent dimer are visualized by a cyan surface.

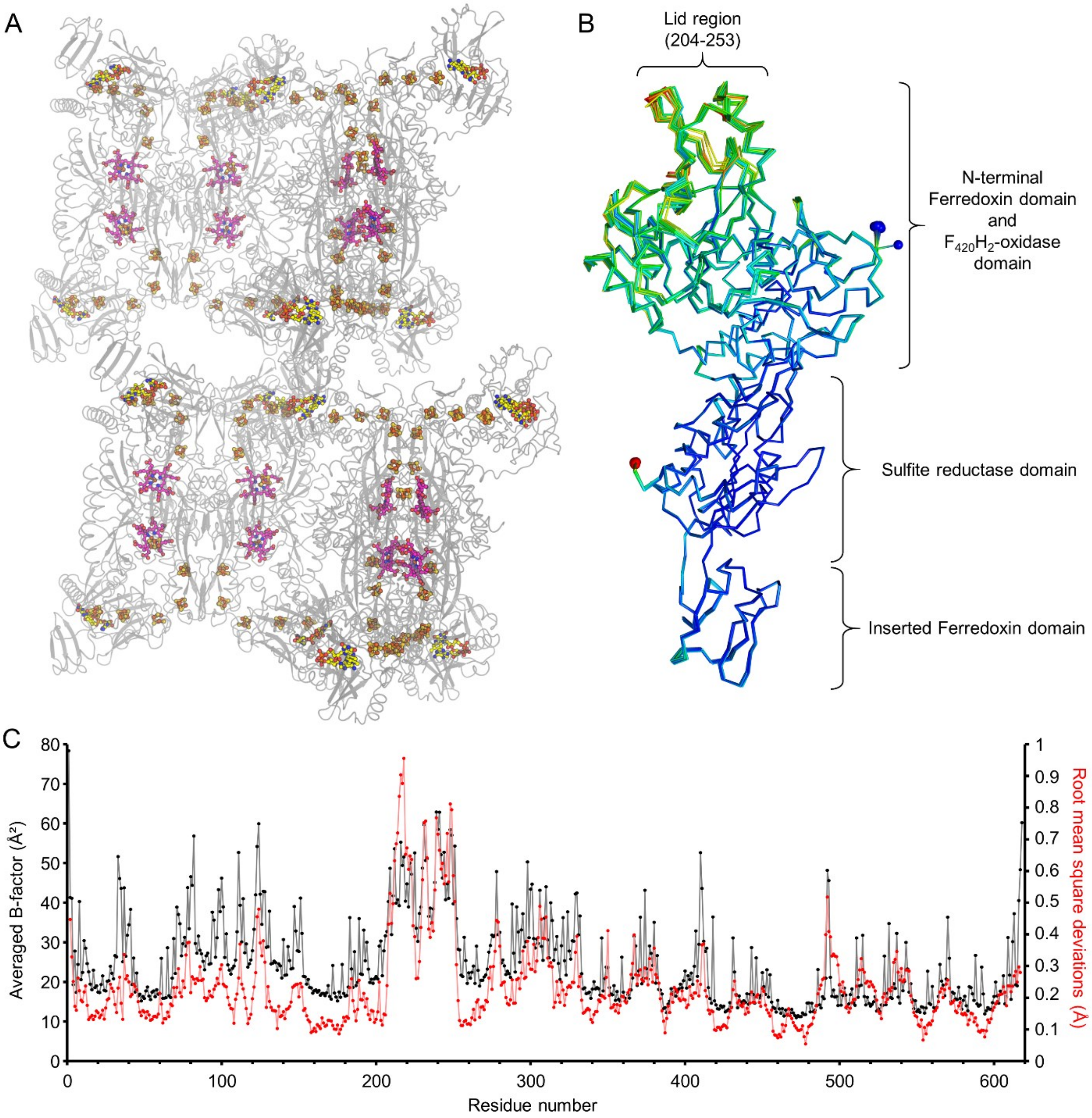


**Supplementary Figure S6.** Asymmetric unit content and B-factor profile of *Mt*Fsr. (A) Representation of the four homotetramers contained in the asymmetric unit of *Mt*Fsr emphasizing the 96 [4Fe-4S]-clusters, the 16 FADs (in yellow) and 16 sirohemes (in pink) shown in balls and sticks. (B) Superposition of all sixteen chains from the asymmetric unit in *Mt*Fsr. The average rmsd was 0.14 Å with 514-Cα aligned. The N-terminus of each chain is shown by a blue sphere and the C-terminus by a red sphere. The models are colored according to their B-factors values; blue to red indicates low to high B-factors, respectively. (C) B-factor profile (in black) for each residue of *Mt*Fsr, values are averaged from the 16 chains composing the asymmetric unit. On the same graph is overlaid the averaged root mean square deviations (rmsd, in red) of the corresponding Cα. Each chain, composing the asymmetric unit, has been superposed on chain A by the software superpose (11) and rmsds were averaged and plotted.

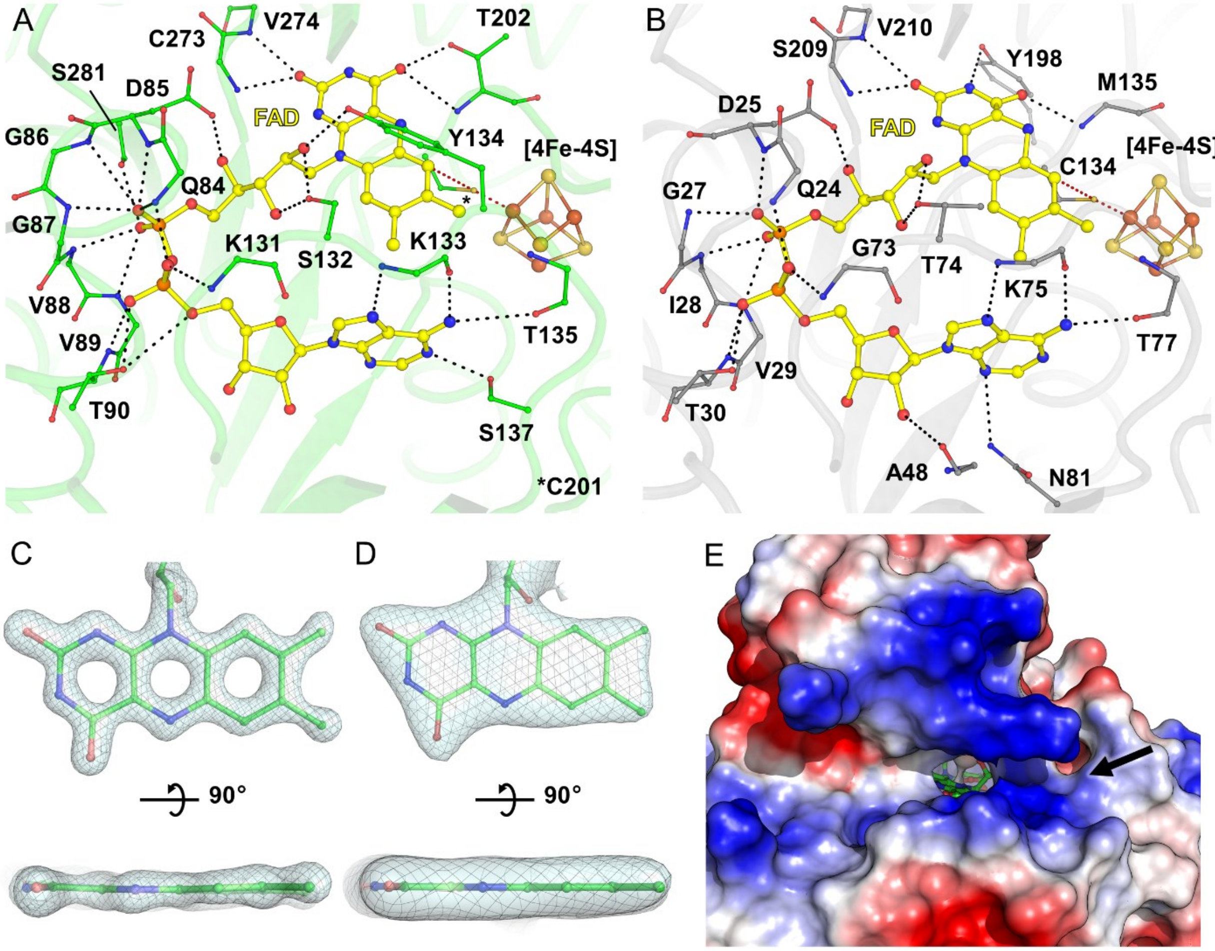


**Supplementary Figure S7.** Active site of the $F_{420}H_2$-oxidase domain. (A) Close-up of the FAD binding site in $F_{420}H_2$-oxidase domain from *Mt*Fsr. (B) FAD binding in FrhB from *M. marburgensis* (PDB 4OMF). In A and B, the residues binding the FAD are represented in balls and sticks. Hydrogen bonds involved in FAD binding are shown by black dashes. (C and D) $2F_o$-$F_c$ map for the FAD in *Mt*Fsr (C) and *Mj*Fsr (D) contoured to 3- and 1.5-σ, respectively. The isoalloxazine heterocycle is only slightly bent. (E) Electrostatic charge profile around the $F_{420}H_2$-oxidase active site in *Mt*Fsr. Acidic to basic patches on the surface are colored in red and blue, respectively. The $F_{420}H_2$ is suspected to bind to the positively charged area (indicated by a black arrow) surrounding the FAD.

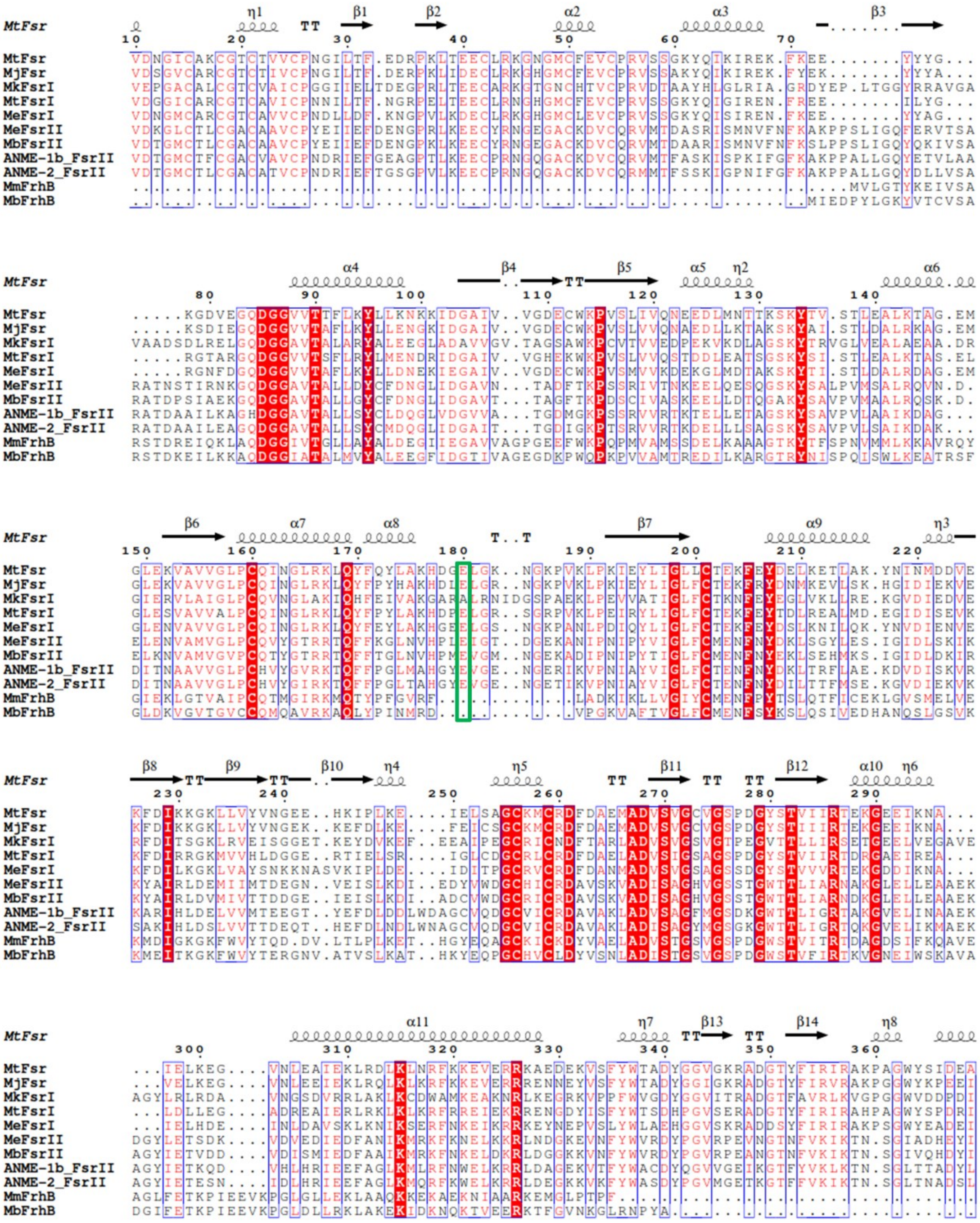


**Supplementary Figure S8.** Sequence conservation across Fsr Group I and II as well as FrhB. Perfectly conserved residues are highlighted with a red background. The glutamate involved in the [4Fe-4S]-cluster 3 binding is shown by a green square. MtFsr: *M. thermolithotrophicus*, MjFsr: *M. jannaschii* (Q58280.1), MkFsrI*: Methanopyrus kandleri* (WP_011019168.1); MtFsrI: *Methanothermobacter marburgensis* (ADL58324.1); MeFsrI: *Methanohalobium evestigatum* (WP_013194157.1); MeFsrII: *Methanohalobium evestigatum* (WP_013194630.1); MbFsrII *Methanococcoides burtonii* (WP_011498749.1); ANME-1b_FsrII: Fsr from Methanophagales archaeon belonging to ANME-1 cluster (RCV63578.1); ANME-2_FsrII, Fsr from ANME-2 cluster archaeon HR1 (PPA79744.1); MmFrhB: *Methanothermobacter marburgensis* (ADL59254.1); MbFrhB *Methanosarcina barkeri* (WP_048177139.1). Sequence alignment was done using Clustal Omega (12), secondary structure prediction was performed with ESPript 3.0 (13).

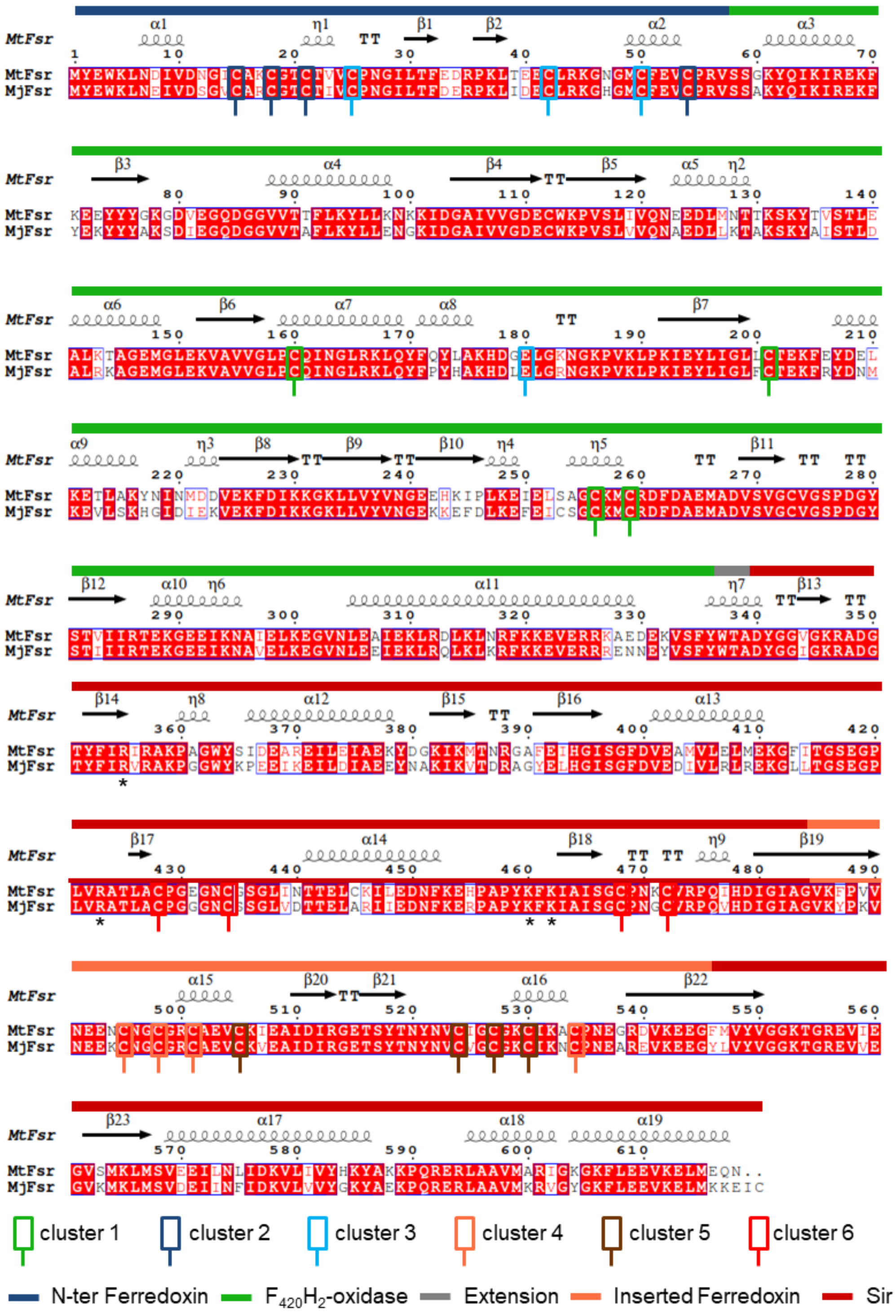

**Supplementary Figure S9.** Sequence alignment between *Mt*Fsr and *Mj*Fsr. Cysteines and the glutamate involved in direct [4Fe-4S]-cluster binding are highlighted, as well as the different domains in Fsr. Sequence alignment was done using Clustal Omega (12), secondary structure prediction was performed with ESPript 3.0 (13). Cluster 6 corresponds to the electronically coupled siroheme–[4Fe–4S] cluster. The stars (*) indicate residues near the siroheme proposed to bind $SO_3^{2-}$.

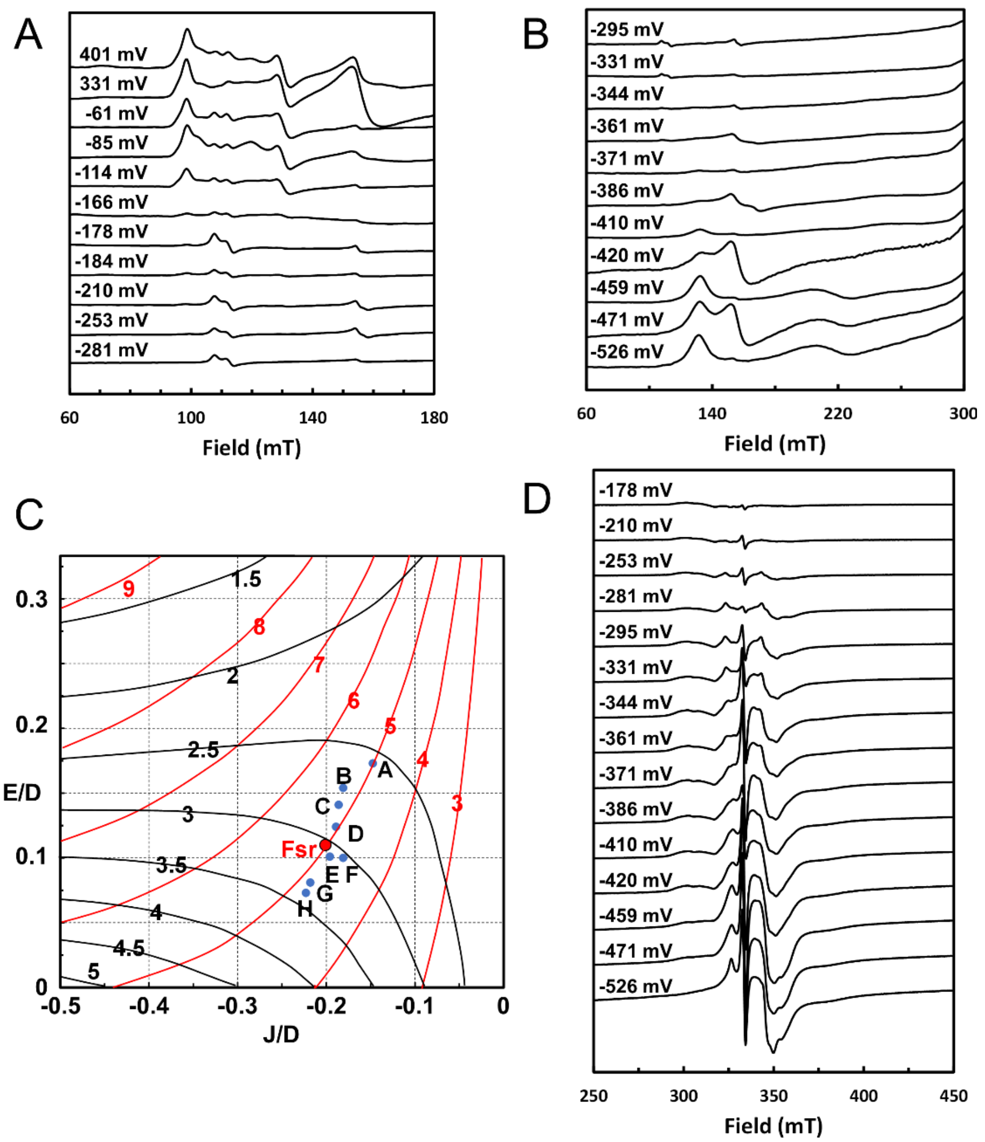


**Supplementary Figure S10.** EPR spectra of the dye-mediated redox titrations of *Mt*Fsr and *g*-values as function of J/D and E/D as described by J. A. Christner et al. 1984. (A, B and D) The redox potentials at which samples were frozen are indicated. EPR intensities were scaled to correct for differences in concentration. EPR conditions: temperature, 10 K; modulation frequency, 100 kHz; modulation amplitude, 1.0 mT; microwave frequency 9.353 GHz; microwave power 20 mW (panel (D) 0.2 mW). (C) Contours of the two highest *g*-values of the coupled ferrous siroheme-[4Fe-4S]$^{1+}$ system as function of J/D and E/D according to Fig. 5 from (14). The blue points are from *E. coli* sulfite reductase: A and B, KCl (two species); C, KF or KBr; E, urea; F, sodium formate; G, $(Gdm)_2SO_4$; H, KBr; D, spinach nitrite reductase; *Mt*FsR is shows as red point.

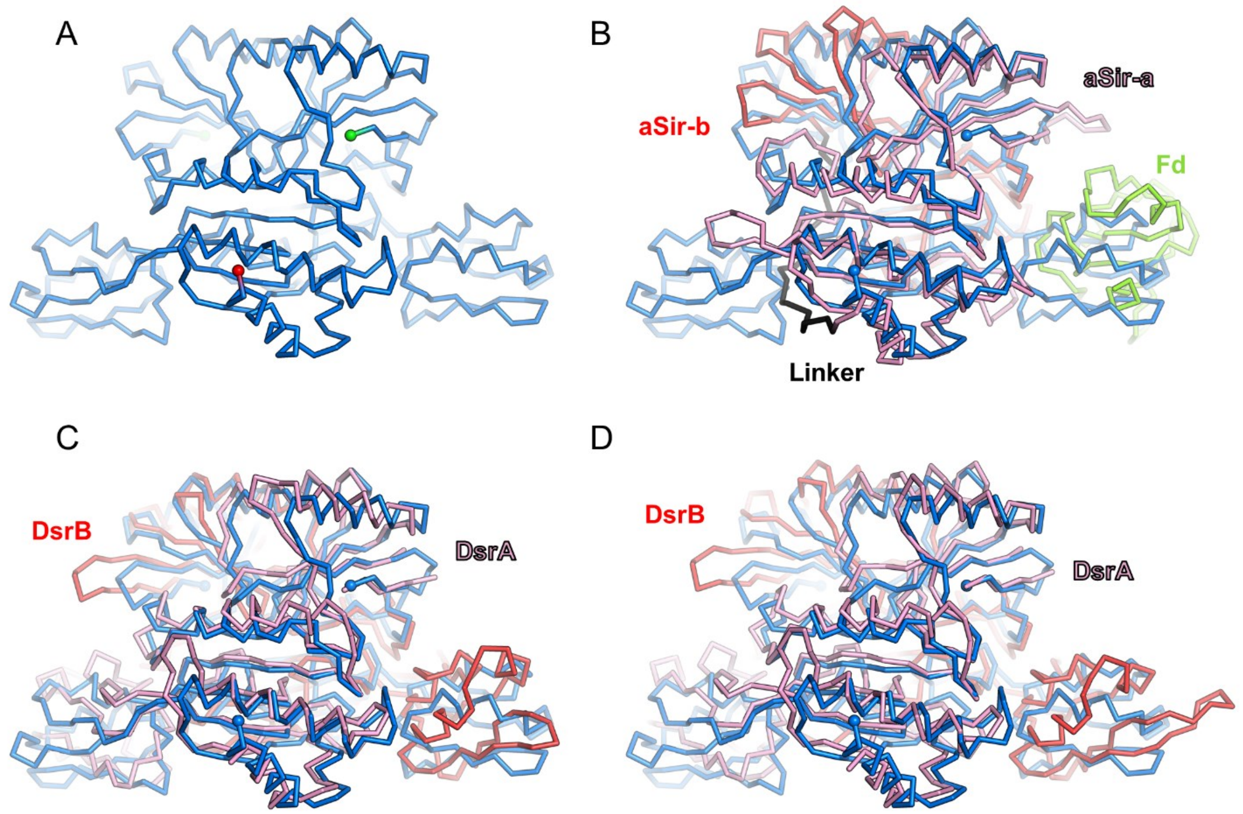


**Supplementary Figure S11.** Fsr shares the common fold of sulfite reductases. (A) The dimeric sulfite reductase domains of *Mt*Fsr with its inserted ferredoxin domains are represented in blue ribbon. N- and C-terminus of the Sir domain are in green and red spheres, respectively. (B) Overall superposition of dimeric *Mt*Fsr (blue) with the aSir from *Zea mays* (PDB 5H92, whole chain rmsd=2.434 Å for 142-Cα aligned and a rmsd=0.964 Å for the most conserved region with 47-Cα aligned). The aSir-a part from *Zea mays* is colored in light pink, and the aSir-b part is colored in red. (C) Overall superposition of dimeric *Mt*Fsr with DsrAB (DsrA in pink, DsrB in red) from *Archaeoglobus fulgidus* (PDB 3MM5, whole chain rmsd=4.152 Å for 225-Cα aligned and an rmsd=0.921 Å with 53-Cα aligned for the most conserved region on one *Mt*Fsr monomer). (D) Overall superposition of dimeric *Mt*Fsr with DsrAB (DsrA in pink, DsrB in red) from *Desulfovibrio vulgaris* (PDB 2V4J, whole chain rmsd=2.819 Å for 157-Cα aligned and a rmsd=0.961 Å with 51-Cα aligned for the most conserved region on one *Mt*Fsr monomer). (B-D) The extensions contained in aSir and DsrAB which are not common to Fsr have been removed for clarity.

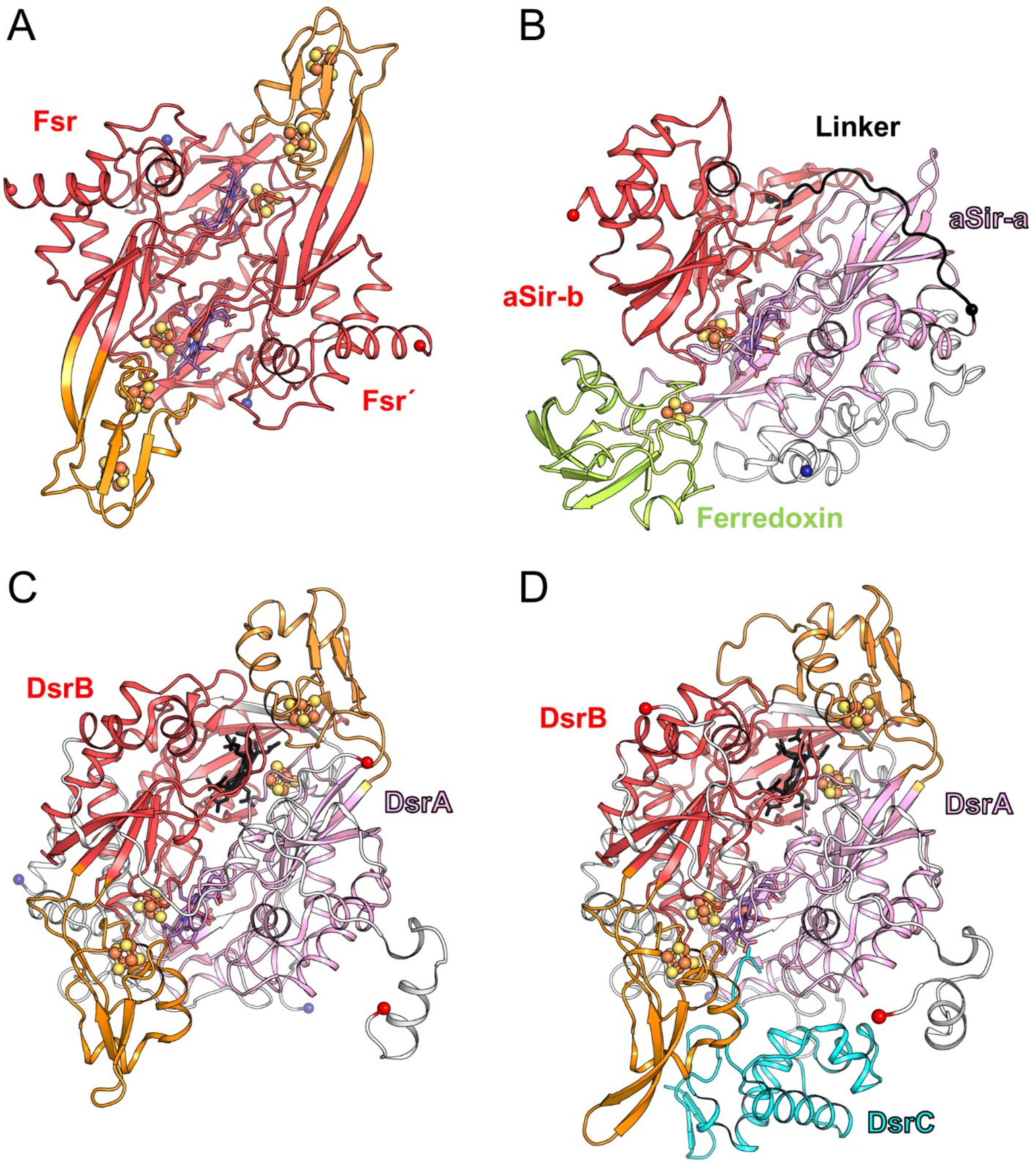


**Supplementary Figure S12.** Overall structural comparison between Fsr, aSir and dSir. Cut-through view shown in cartoon of one dimer for Fsr and DsrAB. Ligands are shown as balls and sticks. (A) Sulfite reductase part with the inserted ferredoxin domain of *Mt*Fsr. Fsr´ corresponds to the opposite monomer. (B) aSir from *Zea mays* and its [2Fe-2S]-ferredoxin colored in light green (PDB 5H92). (C) DsrAB from *A. fulgidus* (PDB 3MM5) and (D) DsrABC from *D. vulgaris* (PDB 2V4J). The inserted ferredoxin domains of Fsr, DsrA and DsrB are colored in orange. The catalytic siroheme in DsrAB is colored in deep purple and the structural siroheme is colored in black. DsrAB from *D. vulgaris* contains sirohydrochlorin instead of a siroheme.

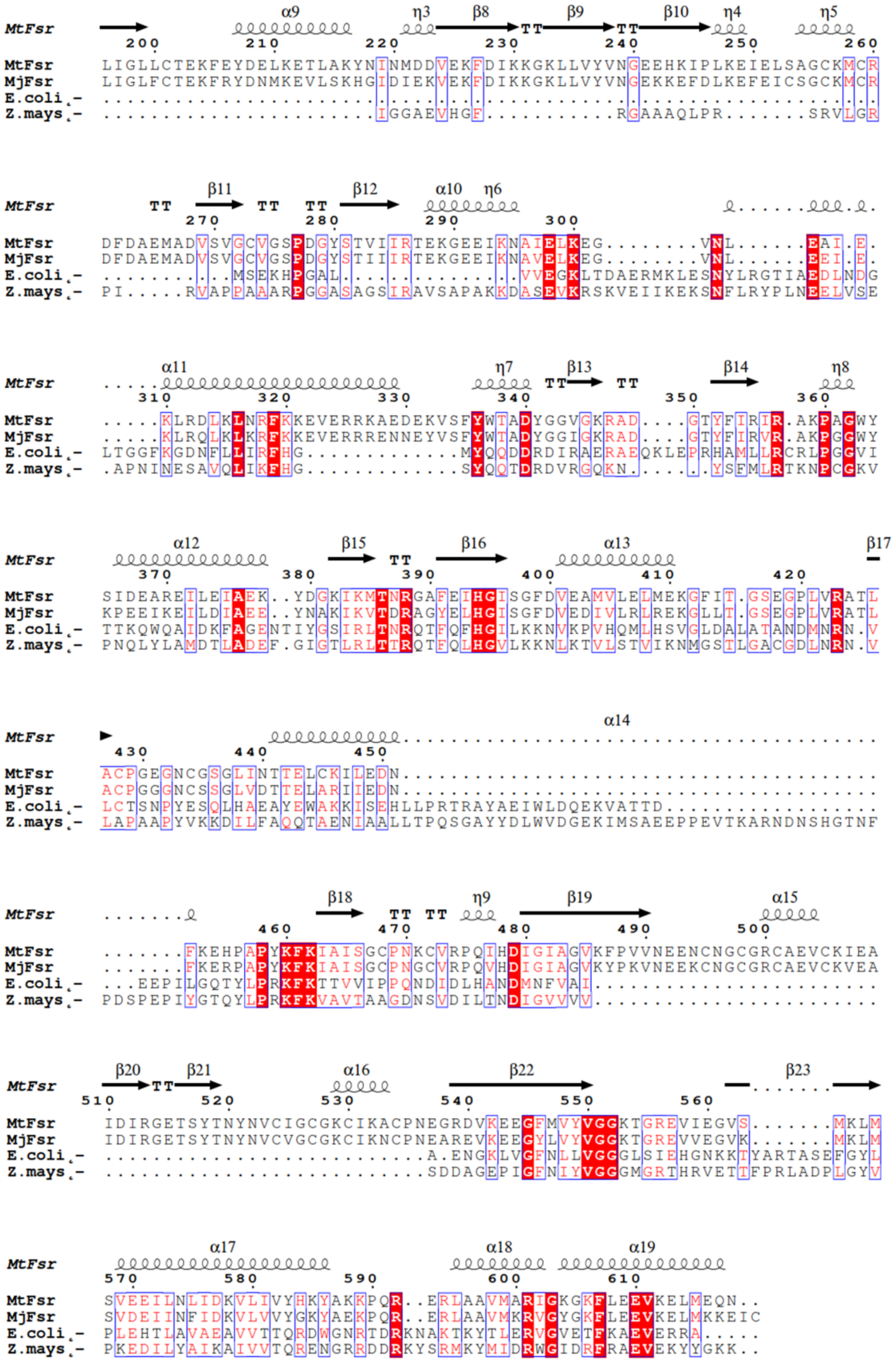

**Supplementary Figure S13.** Sequence conservation across the C-terminal half of Fsr (*Mt*Fsr: 339-618) and aSir-a. Perfectly conserved residues are highlighted with a red background. MtFsr: *M. thermolithotrophicus*, MjFsr: *M. jannaschii*. Z.mays*: Zea mays* (PDB 5H92, for residues: 1-392); E.coli: *Escherichia coli* (PDB 2GEP, for residues: 1-327). Sequence alignment was done using Clustal Omega (12), secondary structure prediction was performed with ESPript 3.0 (13).

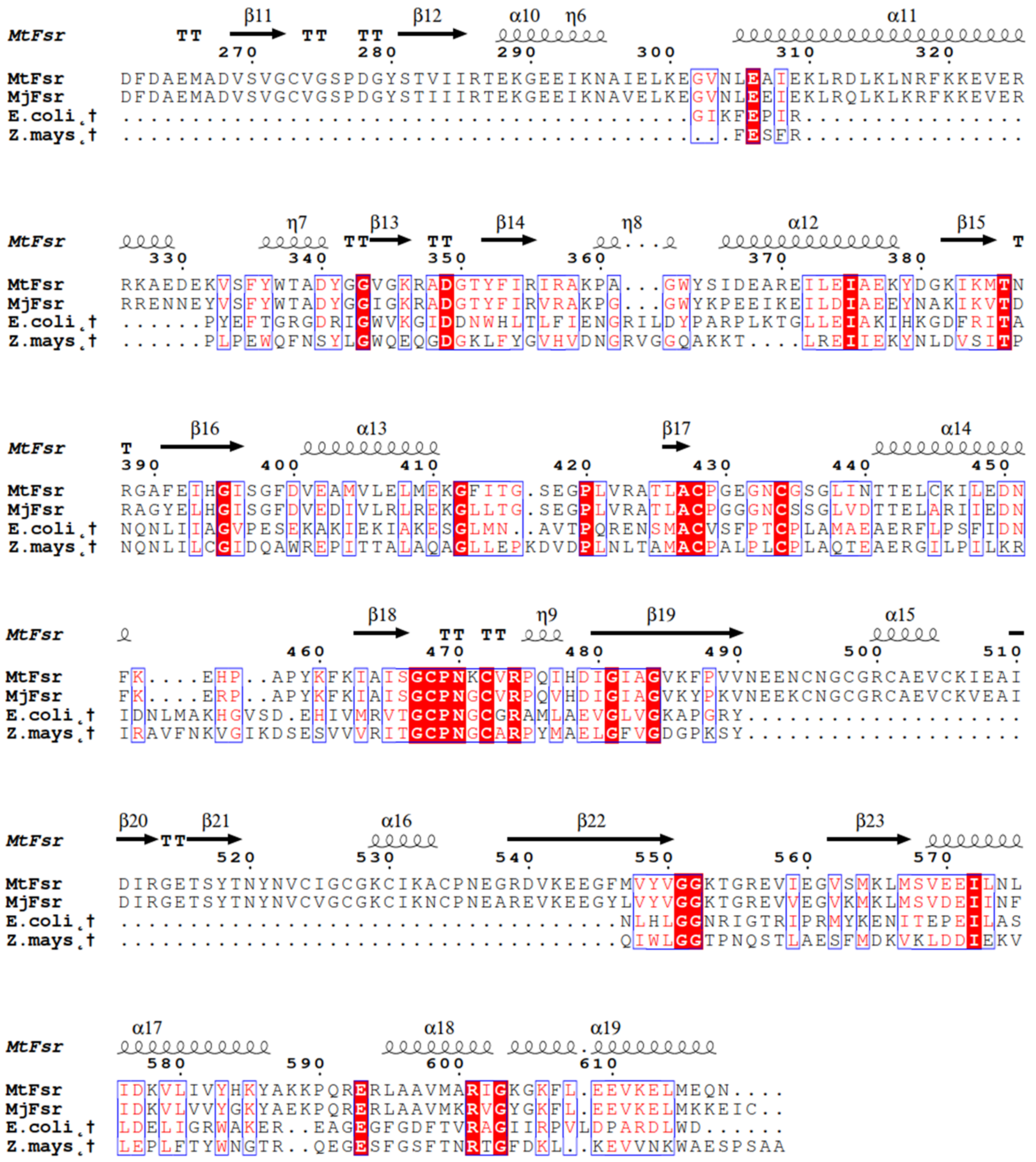

**Supplementary Figure S14.** Sequence conservation across the C-terminal half of Fsr (*Mt*Fsr: 339-618) and aSir-b. Perfectly conserved residues are highlighted with a red background. MtFsr: *M. thermolithotrophicus*, MjFsr: *M. jannaschii*. Z. mays*: Zea mays* (PDB 5H92, used residues: 393-653); E.coli: *Escherichia coli* (PDB 2GEP, shown residues: 328-570). Sequence alignment was done using Clustal Omega (12), secondary structure prediction was performed with ESPript 3.0 (13).

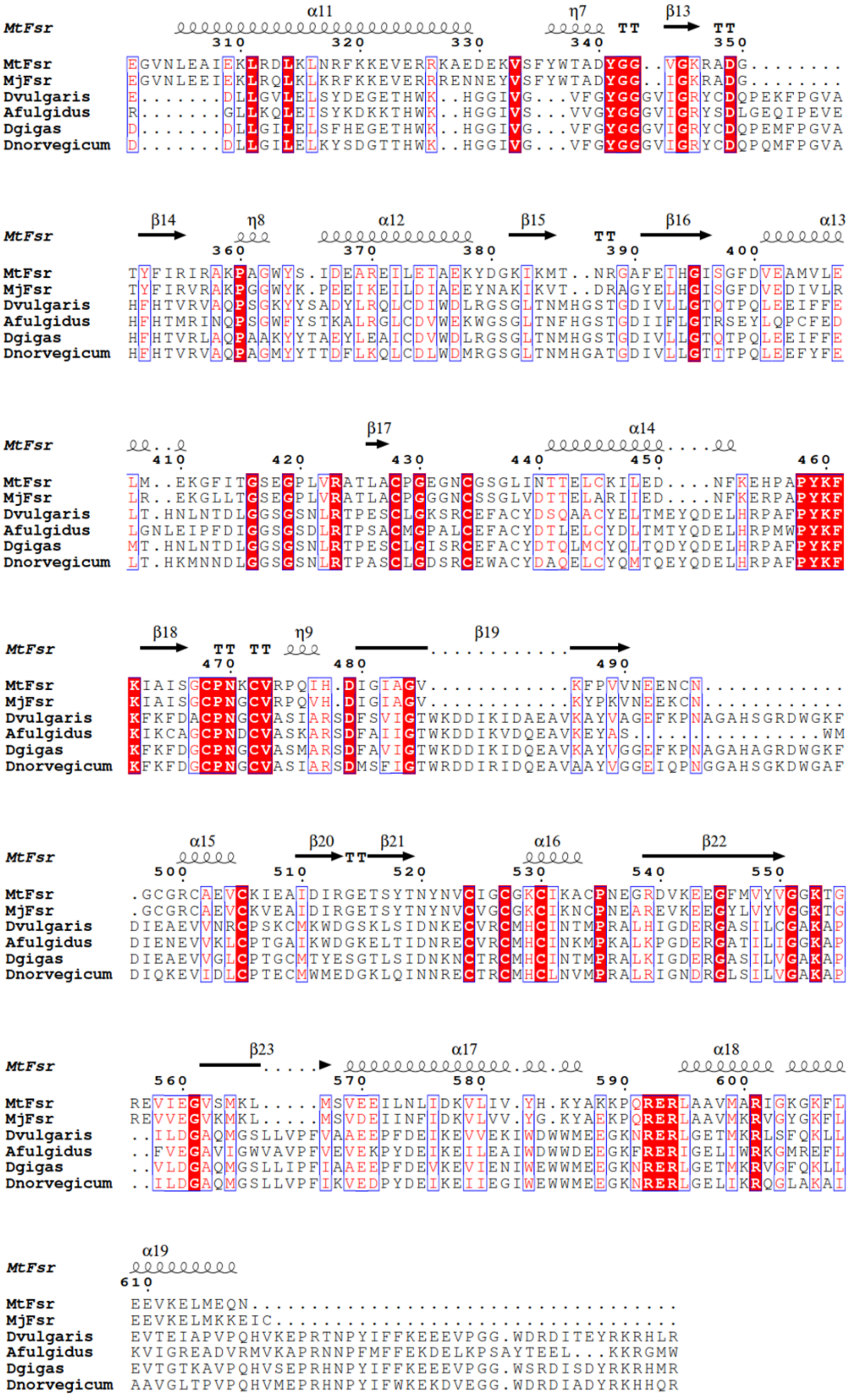

**Supplementary Figure S15.** Sequence conservation across the C-terminal half of Fsr (*Mt*Fsr: 339-618) and DsrA. Perfectly conserved residues are highlighted with a red background. MtFsr: *M. thermolithotrophicus*, MjFsr: *M. jannaschii*. Dvulgaris*: D. vulgaris* (PDB 2V4J); Afulgidus: *A. fulgidus* (PDB 3MM5); Dgigas: *Desulfovibrio gigas* (PDB 3OR1); Dnorvegicum: *Desulfomicrobium norvegicum* (PDB 2XSJ). Sequence alignment was done using Clustal Omega (12), secondary structure prediction was performed with ESPript 3.0 (13).

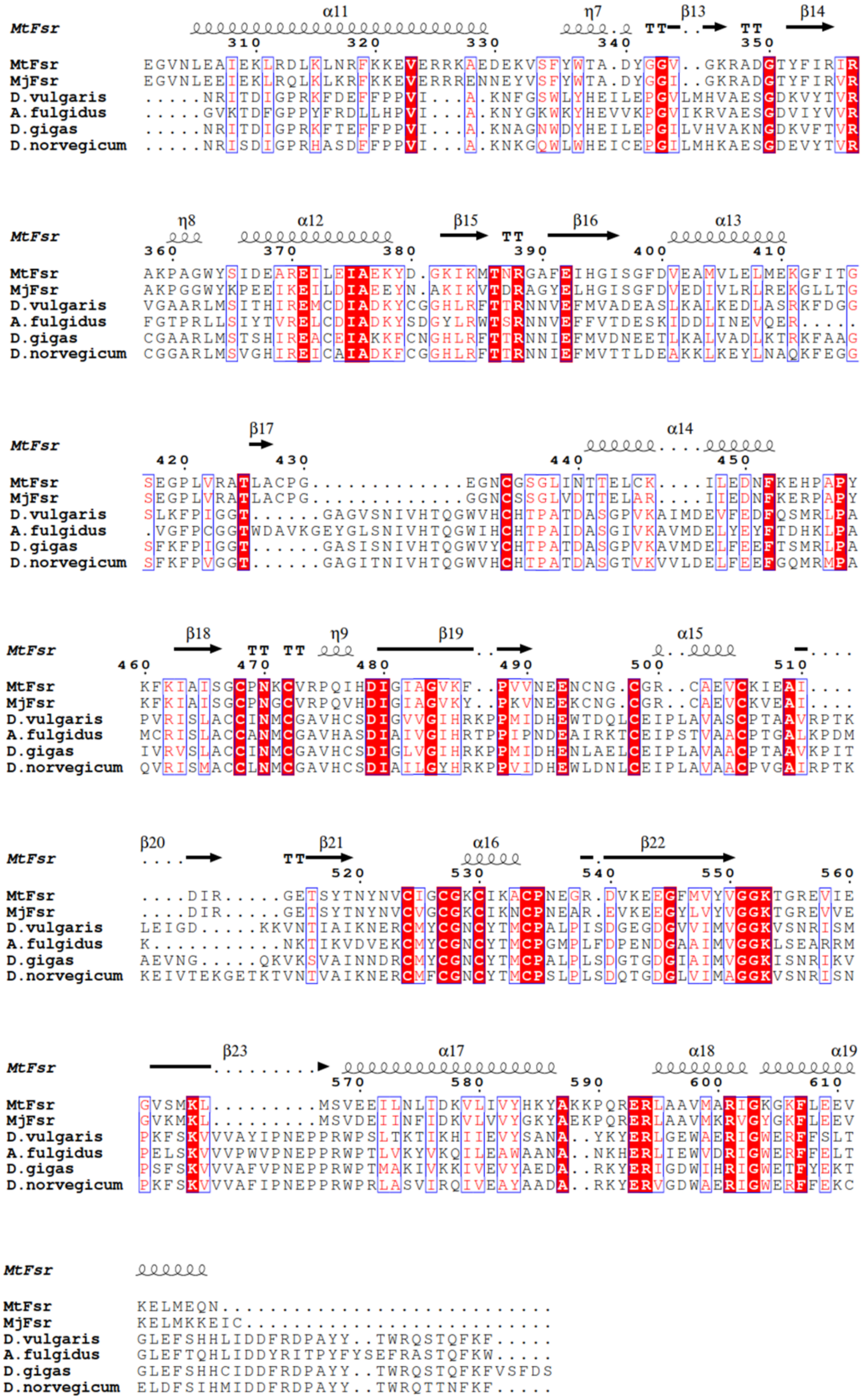


**Supplementary Figure S16.** Sequence conservation across the C-terminal half of Fsr (*Mt*Fsr: 339-618) and DsrB. Perfectly conserved residues are highlighted with a red background. MtFsr: *M. thermolithotrophicus*, MjFsr: *M. jannaschii*. Dvulgaris*: Desulfovibrio vulgaris* (PDB 2V4J); Afulgidus: *Archaeoglobus fulgidus* (PDB 3MM5); Dgigas: *Desulfovibrio gigas* (PDB 3OR1); Dnorvegicum: *Desulfomicrobium norvegicum* (PDB 2XSJ). Sequence alignment was done using Clustal Omega (12), secondary structure prediction was performed with ESPript 3.0 (13).

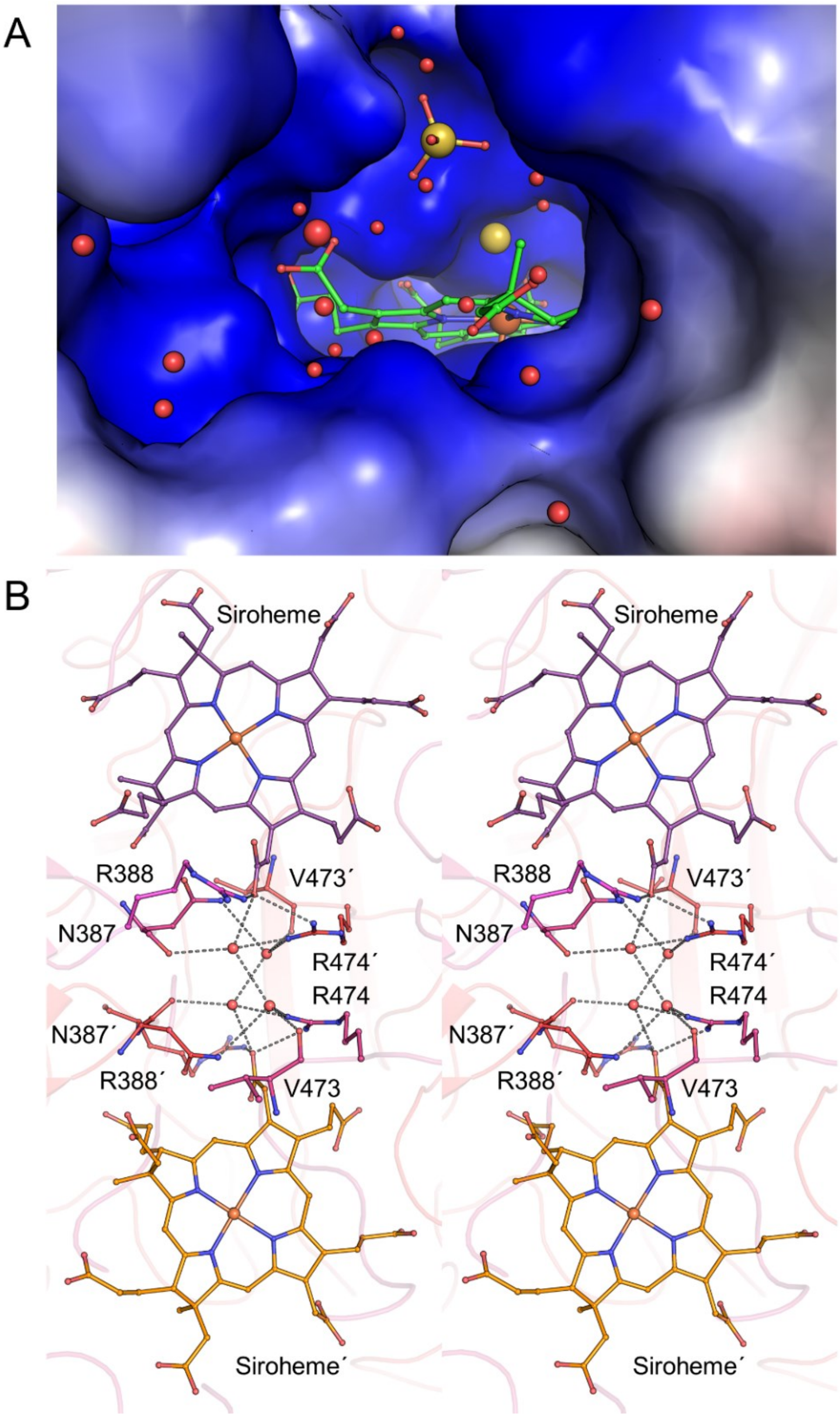


**Supplementary Figure S17.** Siroheme position within *Mt*Fsr (A) Sirohemes in Fsr are accessible via a positively charged solvent channel. Carbon, oxygen, nitrogen, sulfur and iron are colored in green, red, blue, yellow and orange, respectively. Electrostatic charge profile shown in surface is colored in red and blue to represent acidic and basic patches, respectively. In *Mt*Fsr, a sulfate molecule sits next to the active site. (B) Stereo view of the intra-dimeric sirohemes in *Mt*Fsr, in which each chain is differently colored. Primed labels indicate residues belonging to the dimeric partner. Sirohemes, waters and residues involved in the channel are represented as balls and sticks. The distance between the two closest siroheme carboxylate groups is 9.4 Å. This close contact would theoretically allow an internal electron transfer between both sirohemes.

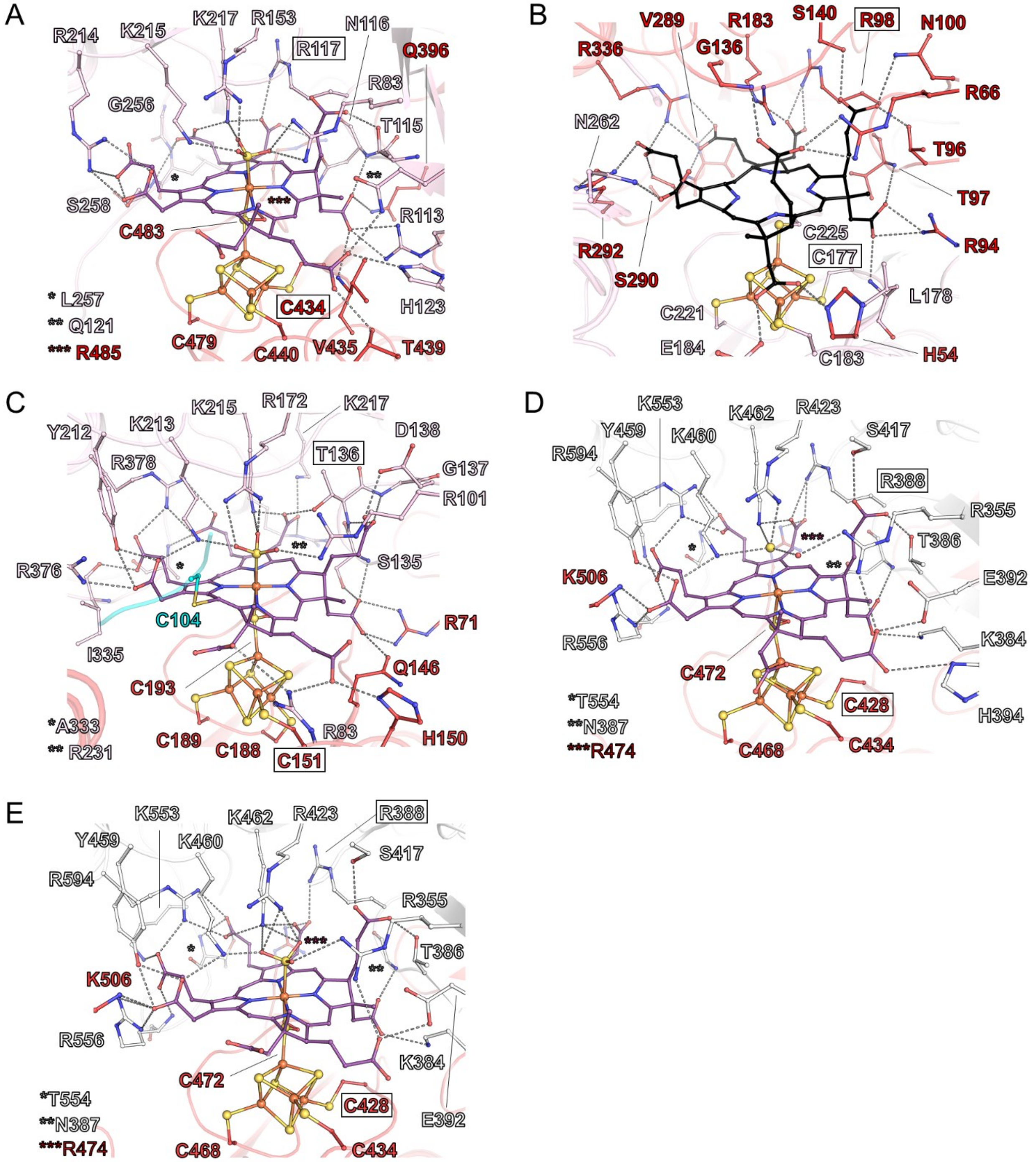


**Supplementary Figure S18.** Siroheme coordination in sulfite reductases. (A) Siroheme site in aSir from *E.coli* (PDB 1AOP). aSir-a and aSir-b parts are colored in light pink and red, respectively. (B) Structural active site in dSir from *D. vulgaris*, containing a sirohydrochlorin (in black) (PDB 2V4J, DsrA is colored in light pink and DsrB in red). (C) Functional active site in dSir from *D. vulgaris*. DsrC is shown in cyan with its cysteine interacting with the siroheme. (D) Siroheme coordination in *Mt*Fsr, where a $S^{2-}$ ion was modelled in the active site, whereas (E) in *Mj*Fsr a $SO_3^{2-}$ ion was tentatively modelled in the active site. Each monomer is differently colored (white and red). (A-E) Hydrogen bonds and salt-bridges involved in the coordination of the siroheme and its axial ligand are shown as black dashes. Residues involved in these interactions are in balls and sticks. Framed residues are the most important difference between aSirs and dSirs.

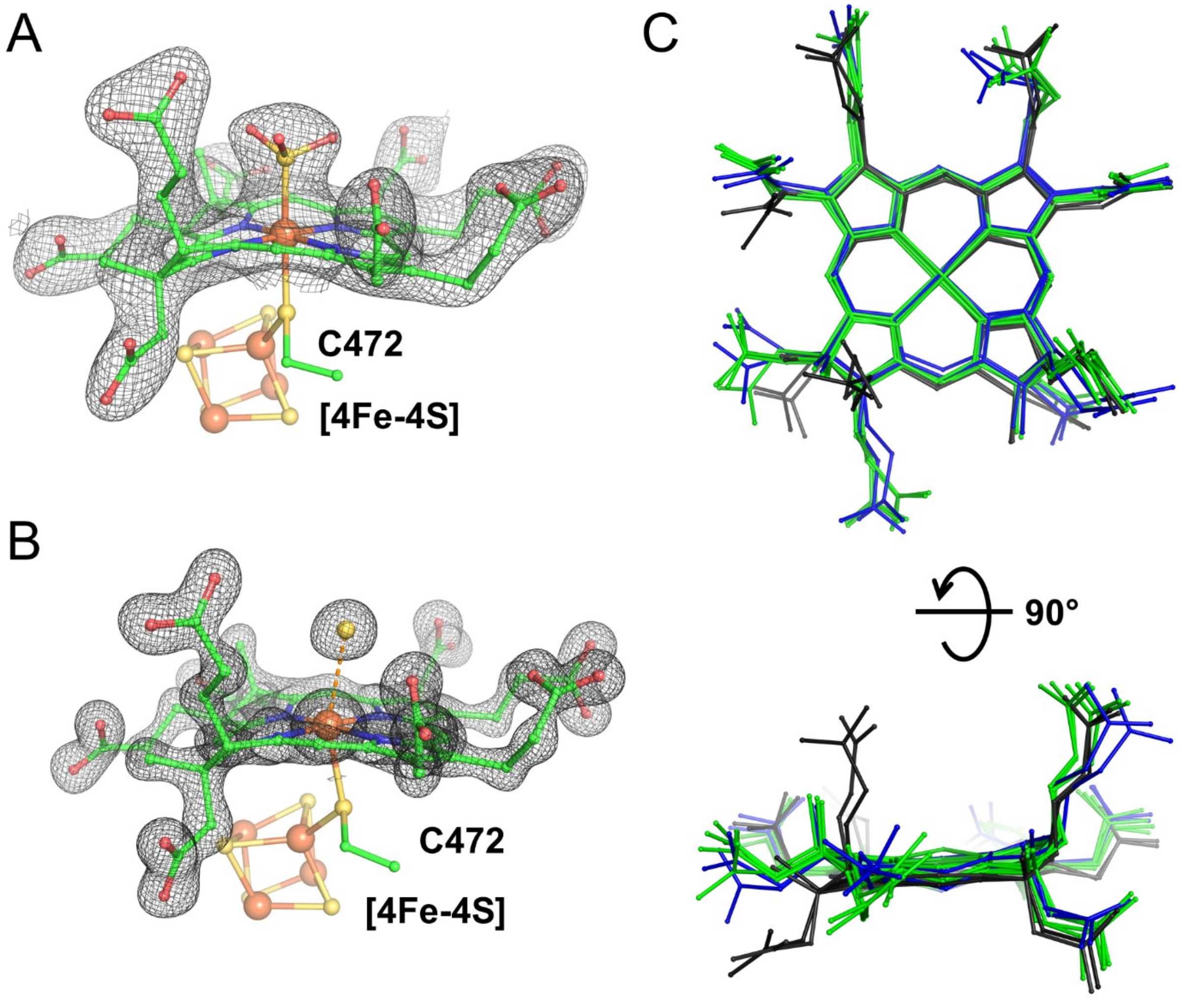


**Supplementary Figure S19.** Siroheme axial ligands and conformation. (A-B) Close up of the axial ligands bound on the siroheme from *Mj*Fsr (A) and *Mt*Fsr (B). The 2$F_o$-$F_c$ map of the siroheme and $SO_3^{2-}$ is contoured to 1.5-σ in *Mj*Fsr. The 2$F_o$-$F_c$ map of the siroheme and $S^{2-}$ is contoured to 3-σ in *Mt*Fsr. In *Mj*Fsr the Fe-siroheme is equidistant (2.3 Å) to the sulfur from the modelled $SO_3^{2-}$ and the bridging-sulfur of the cysteine 472, suggesting a tight covalent binding. In *Mt*Fsr, the bridging-sulfur of the cysteine 472 is at a distance of 2.6 Å to the Fe-siroheme and the sulfur from the modelled $S^{2-}$ is 2.9 Å distant to the Fe-siroheme, indicating a loose binding of the $S^{2-}$, which might result from a reduction event by X-ray radiation (15). (C) Siroheme superposition between aSirs (1AOP, 5H92), dSirs (3MM5, 2V4J) and Fsr. Siroheme from aSirs and Fsr are colored in green, structural siroheme/sirohydrochlorin from dSirs are shown in black, functional sirohemes from dSirs are shown in blue.

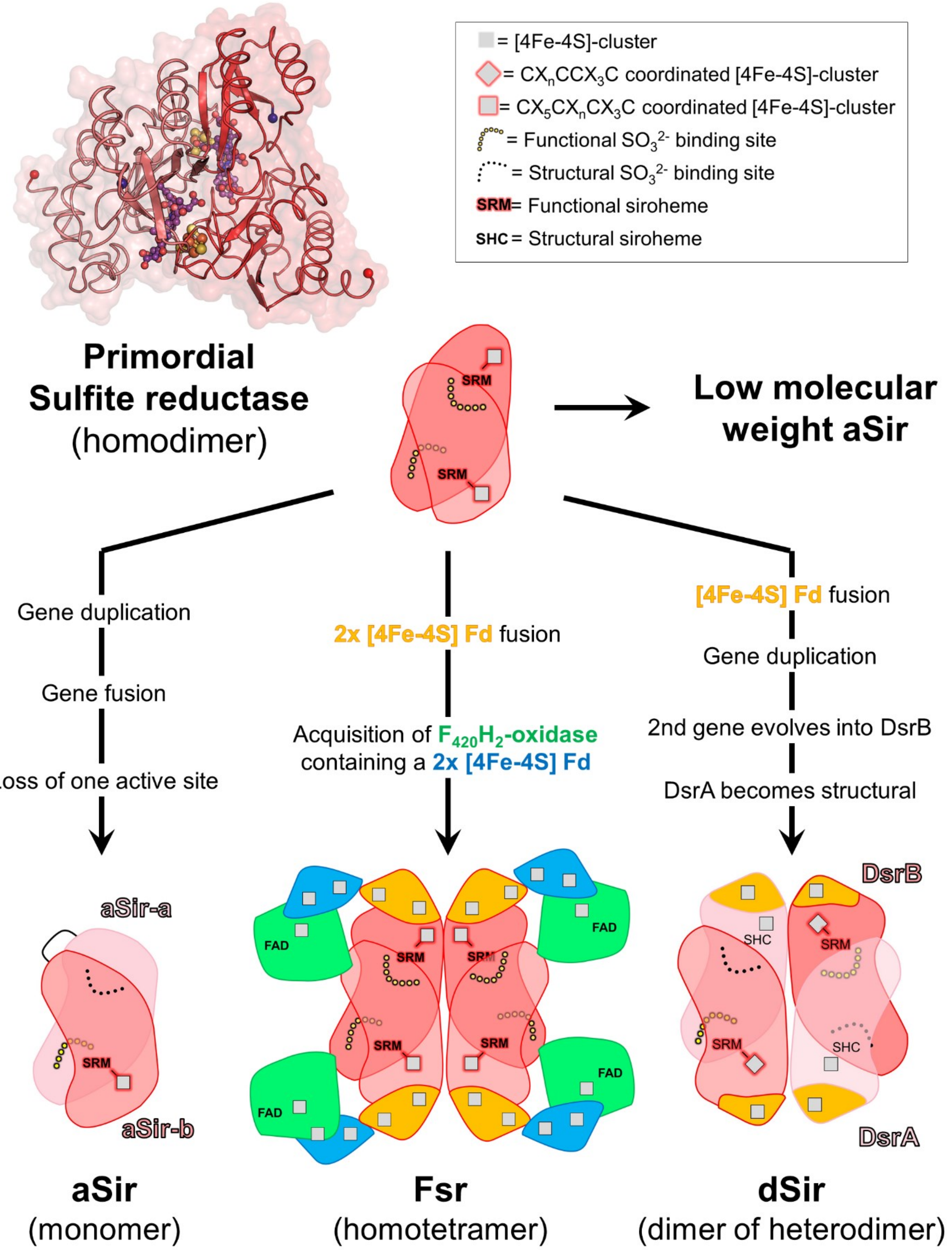


**Supplementary Figure S20.** Theoretical evolutionary scenario of sulfite reductases. The proposed route is based on the assumption that aSir, dSir and Fsr could have evolved from a comon ancestor. The primordial sulfite reductase model corresponds to the elementary sulfite reductase core of *Mt*Fsr structure.
